# Simultaneous proteome localization and turnover analysis reveals spatiotemporal features of protein homeostasis disruptions

**DOI:** 10.1101/2023.01.04.521821

**Authors:** Jordan Currie, Vyshnavi Manda, Sean K. Robinson, Celine Lai, Vertica Agnihotri, Veronica Hidalgo, R. W. Ludwig, Kai Zhang, Jay Pavelka, Zhao V. Wang, June-Wha Rhee, Maggie P. Y. Lam, Edward Lau

## Abstract

The functions of proteins depend on their spatial and temporal distributions, which are not directly measured by static protein abundance. Under endoplasmic reticulum (ER) stress, the unfolded protein response (UPR) pathway remediates proteostasis in part by altering the turnover kinetics and spatial distribution of proteins. A global view of these spatiotemporal changes has yet to emerge and it is unknown how they affect different cellular compartments and pathways. Here we describe a mass spectrometry-based proteomics strategy and data analysis pipeline, termed Simultaneous Proteome Localization and Turnover (SPLAT), to measure concurrently the changes in protein turnover and subcellular distribution in the same experiment. Investigating two common UPR models of thapsigargin and tunicamycin challenge in human AC16 cells, we find that the changes in protein turnover kinetics during UPR varies across subcellular localizations, with overall slowdown but an acceleration in endoplasmic reticulum and Golgi proteins involved in stress response. In parallel, the spatial proteomics component of the experiment revealed an externalization of amino acid transporters and ion channels under UPR, as well as the migration of RNA-binding proteins toward an endosome co-sedimenting compartment. The SPLAT experimental design classifies heavy and light SILAC labeled proteins separately, allowing the observation of differential localization of new and old protein pools and capturing a partition of newly synthesized EGFR and ITGAV to the ER under stress that suggests protein trafficking disruptions. Finally, application of SPLAT toward human induced pluripotent stem cell derived cardiomyocytes (iPSC-CM) exposed to the cancer drug carfilzomib, identified a selective disruption of proteostasis in sarcomeric proteins as a potential mechanism of carfilzomib-mediated cardiotoxicity. Taken together, this study provides a global view into the spatiotemporal dynamics of human cardiac cells and demonstrates a method for inferring the coordinations between spatial and temporal proteome regulations in stress and drug response.

## Introduction

Protein turnover is an important cellular process that maintains the quality and quantity of protein pools in homeostasis, and involves fine regulations of the rates of synthesis and degradation of individual proteins. A close relationship exists between turnover kinetics with the spatial distribution of proteins. Cellular organelles including the cytosol, endoplasmic reticulum (ER), and mitochondria are equipped with distinct quality control and proteolytic mechanisms that maintain protein folding and regulate protein degradation in a localization dependent manner (Lemberg and Strisovsky, 2021; Mårtensson et al., 2019; Tsai et al., 2022). Newly synthesized proteins need to be properly folded and trafficked to their intended subcellular localization through subcellular targeting and sorting mechanisms, whose capacity has to be coordinated to match the rate of protein synthesis (Chartron et al., 2016; Jan et al., 2014; Lakkaraju et al., 2008). A primary subcellular trafficking mechanism of new proteins is the ER vesicular transport and secretory pathway through the endomembrane system, which is a rate-limiting step in the production of membrane and extracellular proteins. A mismatch between temporal synthesis rate and spatial localization capacity can lead to ER stress and subsequently mislocalization of newly synthesized proteins (Hetz et al., 2020).

Disruption of protein turnover and homeostasis is broadly implicated in human diseases including cardiomyopathies, cancer, and neurodegeneration (Hetz et al., 2020; Ren et al., 2021). In stressed cells, the accumulation of misfolded proteins triggers the unfolded protein response (UPR), which involves signaling pathways that suppress protein synthesis and promote protein folding and proteolysis to relieve proteostatic stress. At the same time, UPR activation invokes a spatial reorganization of the proteome, including but not limited to the transient translocation of UPR pathway mediators to the nucleus during initial stress response, the retrotranslocation of misfolded ER proteins to the cytosol for proteasomal clearance under endoplasmic-reticulum-associated protein degradation (ERAD), and the sequestration of RNA and RNA-binding proteins into stress granules. Despite ongoing research, how the cellular proteome reorganizes under proteostatic stress remains incompletely understood, and the full scope of differentially localized proteins and pathways remains to be elucidated.

We wonder how the spatial and temporal regulations of proteins change in conjunction under UPR, such as whether proteins with differential spatial distributions are also differentially turned over. Advances in mass spectrometry methods have allowed the turnover rate and subcellular localization of proteins to be measured on a large scale. The turnover rate and half-life of proteins can be measured using stable isotope labeling in cells and in intact animals followed by mass spectrometry measurements of isotope signatures and kinetics modeling to derive rate constants (Claydon and Beynon, 2012; Doherty et al., 2009; Hammond et al., 2022; Schwanhäusser et al., 2011). Quantitative comparison of turnover rates provides a temporal view into proteostatic regulations and can implicate new pathological signatures and pathways over steady-state mRNA and protein levels (Andrews et al., 2022; Lam et al., 2014; Lau et al., 2018). In parallel, spatial proteomics methods have allowed increasing power to discern the subcellular localization of proteins on a large scale (Christopher et al., 2021; Geladaki et al., 2019; Kennedy et al., 2020; Mulvey et al., 2021; Orre et al., 2019). In recent work using a differential solubility fractionation strategy and mass spectrometry, we observed broad substantial rearrangement of proteins across three subcellular fractions in an acute paraquat challenge model of UPR in the mouse heart, consistent with protein translocation being an important layer of proteome regulation under proteostatic stress (Dostal et al., 2020). Nevertheless, an integrated strategy that can simultaneously measure protein turnover kinetics and spatial information has thus far not been realized.

Here we extended protein turnover measurements to include subcellular localization dynamics, by integrating dynamic SILAC labeling with differential ultracentrifugation-based spatial proteomics profiling strategies. We describe an experimental strategy and computational analysis pipeline to perform <u>s</u>imultaneous <u>p</u>roteome localization <u>a</u>nd turnover (SPLAT) measurements in baseline and stressed cells. SPLAT builds on prior work in protein turnover measurements and subcellular localization profiling, by combining dynamic SILAC isotope labeling, differential ultracentrifugation, isobaric TMT labeling, and kinetic modeling. This strategy allows for concurrent measurement of changes in the turnover dynamics and subcellular distributions of whole cell proteomes under perturbation in a single experiment. Applying this method to human AC16 cardiac cells under thapsigargin- and tunicamycin-induced UPR and to human induced pluripotent stem cell derived-cardiomyocytes (iPSC-CM) under carfilzomib induced proteasome inhibition, we delineated prominent changes in the spatial and temporal distributions of proteins on a proteome scale. The inclusion of spatial information of light and heavy SILAC labeled proteins moreover allowed disaggregation of the localization and trafficking of new and old protein pools.

## Results

### Simultaneous acquisition of turnover and spatial information using a double labeling strategy

We reason that we can use a hyperplexing strategy to simultaneously encode temporal and spatial protein information through isotope labels in the MS1 and MS2 levels, respectively. Hence, we designed a workflow that combines dynamic SILAC metabolic labeling in cultured cells, with TMT labeling of spatially separated fractions to simultaneously measure new protein synthesis as well as subcellular localization under baseline and perturbation conditions (**Figure 1A**). To determine the rate of protein turnover during control, thapsigargin, and tunicamycin conditions, a dynamic SILAC strategy was used to measure the rate of appearance of post-labeling synthesized protein. Briefly, cells were pulsed with a lysine and arginine depleted media supplemented with heavy labeled lysine and arginine concurrently with drug treatment to label post-treatment synthesized proteins and derive fractional synthesis rates through kinetic modeling.

**Figure 1.**
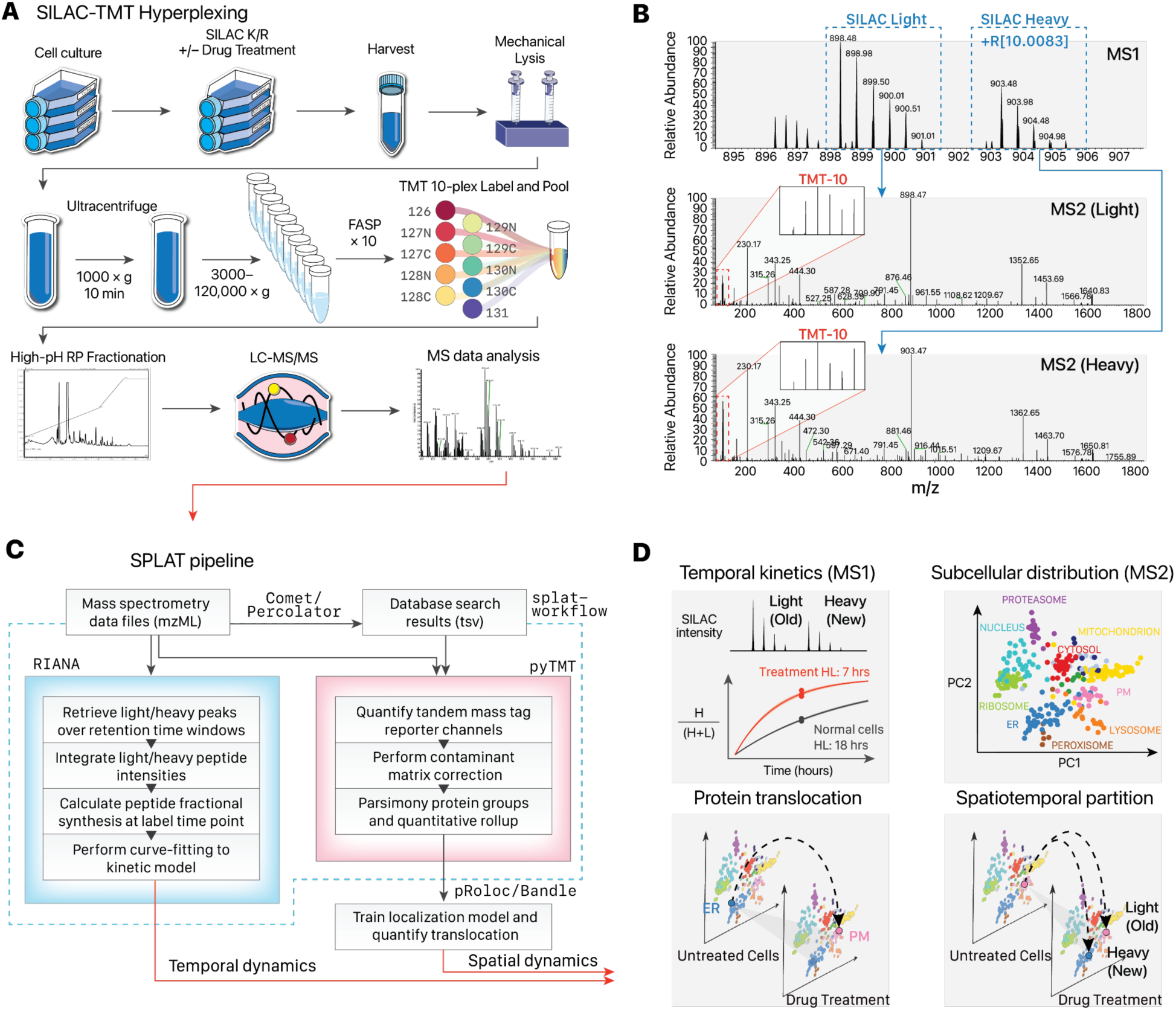
Overview of the SPLAT strategy. **A.** Experimental workflow. Control, thapsigargin-treated, and tunicamycin-treated human A16 cardiomyocytes were labeled with ^13^C6^15^N2 L-Lysine and ^13^C6^15^N4 L-Arginine dynamic SILAC labels. For each condition, 3 biological replicate SPLAT experiments were performed (n=3). After 16 hours, cells were harvested and mechanically disrupted, followed by differential ultracentrifugation steps to pellet proteins across cellular compartments. Proteins from the ultracentrifugation fractions were digested and labeled using tandem mass tag (TMT) followed by mass spectrometry. **B.** Dynamic SILAC labeling allowed differentiation of pre-existing (unlabeled, i.e., SILAC light) and post-labeling (heavy lysine or arginine, i.e., +R[10.0083]) synthesized peptides at 16 hours. The light and heavy peptides were isolated for fragmentation separately to allow the protein sedimentation profiles containing spatial information to be discerned from TMT channel intensities. **C.** Computational workflow. Mass spectrometry raw data were converted to mzML format to identify peptides using a database search engine. The mass spectra and identification output were processed using RIANA (left) to quantitate the time dependent change in SILAC labeling intensities and determine the protein half-life, and using pyTMT (right) to extract and correct TMT channel intensities from each light or heavy peptide MS2 spectrum. The TMT data were further processed using pRoloc/Bandle to predict protein subcellular localization via supervised learning. **D.** Temporal information and spatial information is resolved in MS1 and MS2 levels, respectively. SPLAT allows the subcellular spatial information of the heavy (new) and light (old) subpools of thousands of proteins to be quantified simultaneously in normal and perturbed cells. HL: Half-life.

Upon harvesting, the cells were fractionated to resolve subcellular compartments. We adopted a protein correlation profiling approach. In particular, the LOPIT-DC (Localisation of Organelle Proteins by Isotope Tagging after Differential ultraCentrifugation) method (Geladaki et al., 2019) uses sequential ultracentrifugation to enrich different subcellular fractions from the same samples, which facilitates ease of adoption and reproducibility. Briefly, the cells were lysed under gentle conditions and then sequentially pelleted through ultracentrifugation steps, which pellets subcellular fractions based on their sedimentation rate and which is a function of particle mass, shape, and volume. The ultracentrifugation fractions were each subsequently solubilized, and the extracted proteins were digested and further labeled with tandem mass tag (TMT) isobaric stable isotope labels. The acquired mass spectrometry data therefore carries temporal information in the dynamic SILAC tags and spatial information in the TMT channel intensities (**Figure 1B**).

To process the double isotope encoded mass spectrometry data, we assembled a custom computational pipeline comprising database search and post-processing, and quantification for dynamic SILAC and TMT data (**Figure 1C**). The turnover kinetics information from the dynamic SILAC data is analyzed using a mass spectrometry software tool we previously developed, RIANA (Hammond et al., 2022), which integrates the areas-under-curve of mass isotopomers from peptides over a specified retention time window, then performs kinetic curve-fitting to a mono-exponential model to measure the fractional synthesis rates (FSR) of each dynamic SILAC-labeled (K and R containing) peptide. To extract the ultracentrifugation fraction quantification information for spatial analysis, we developed a new version of the pyTMT tool which we previously described (Dostal et al., 2020), and used it to perform TMT label quantification for the Comet/Percolator workflow. To account for specific challenges related to spatial proteomics data features, we made two new modifications. First, we account for isotope impurities in TMT tags. Because the TMT data are row normalized in the LOPIT-DC design, we incorporated correction of isotope contamination of TMT channels based on the batch contamination data sheet (**Supplemental Table S1**) to account for isotope impurity in fractional abundance calculation from randomized channels across experiments (**Supplemental Methods**). Second, we implemented an isoform-aware quantitative rollup of peptide channel intensities into the protein level for the downstream spatial proteomics analysis. Standard protein inference invokes parsimony rules that assign peptides to the protein within a protein group with the highest level of evidence, but razor peptides can conflate spatial information from different proteins with different localization. Here the TMT-quantified peptides are summed into protein groups using a more conservative aggregate method, such that the identified peptides that are assigned to two or more top-level UniProt protein accessions are discarded to avoid confounding of spatial information in the TMT channels. Moreover, protein groups containing two or more proteins belonging to the same top-level UniProt accession are removed from consideration if one of the non-canonical isoforms contain a unique peptide, and are otherwise rolled up to the canonical protein. Protein isoforms are only included in downstream analysis through quantified isoform-unique peptides. Following RIANA and pyTMT processing, the SPLAT pipeline combines the dynamic SILAC and TMT information by peptides and appends a heavy (“_H”) tag to the UniProt accession of all peptides containing dynamic SILAC modifications for separate localization analyses. The data were then used for temporal kinetics summaries using the MS1 encoded information and subcellular localization classification from the MS2 encoded information (**Figure 1D**). By separately analyzing heavy and light peptides, the subcellular spatial information of the heavy (new) and light (old) subpools of thousands of proteins can be mapped simultaneously in normal and perturbed cells.

### Protein turnover kinetics regulations under unfolded protein response vary by cellular compartments

We applied SPLAT to identify protein spatiotemporal changes in human AC16 cells under UPR induced by 1 µM thapsigargin for 16 hours. Thapsigargin at the dosage and duration used is a common and robust model to induce ER stress and integrated stress response in cardiac and other cell types through the inhibition of sarco/endoplasmic reticulum Ca2+-ATPase (SERCA). Thapsigargin treatment at 16 hours robustly induced known ER stress markers (Glembotski, 2007) including BiP/HSPA5, HSP90B1, PDIA4 (limma FDR adjusted P < 0.01) (**Figure 2A**). Three biological replicate SPLAT experiments were carried out for normal and thapsigargin-treated AC16 cells (n=3 each). We analyzed the spatial fractionation patterns of the proteins following ultracentrifugation and TMT labeling, and classified the subcellular localization of proteins using a Bayesian model BANDLE as previously described (Crook et al., 2018). A spatial classification model is trained separately for each treatment using a basket of canonical organelle markers (**see Methods**) which showed clear separation in PC1 and PC2 in each condition (**Supplemental Figure S1**). The ultracentrifugation profiles of each cellular compartment are highly consistent across treatments and replicates (**Supplemental Figure S2**). To minimize the potential ratio compression that can result from MS2-based TMT quantification, we employed extensive two-dimensional fractionation and narrow isolation window, and verified that identified MS2 spectra had high precursor ion purity (median purity 92–93%) (**Supplemental Figure S3**). We further performed a direct comparison of MS2 and MS3 based quantification on an identical sample (control replicate 2) (**Supplemental Figure S4**), which confirmed that MS2-based quantification produced acceptable spatial resolution, consistent with previous observations (Shin et al., 2020). In total using MS2-based TMT quantification, we mapped the subcellular profiles of 4360 protein features (i.e., 1,820 new and 2,540 old proteins) in normal AC16 cells across 3 biological replicate experiments using a stringent two-peptide filter at 1% FDR of protein identification, with 1946 old proteins and 1,462 new proteins assigned to one of 12 subcellular localization with >95% confidence after removing outliers (see Methods) (**Figure 2B; Supplemental Data S1**). The accuracy of the spatial classification is supported by the observation that 69.5% of assigned proteins in normal AC16 cells contain matching cellular component annotation in Gene Ontology despite the current incompleteness of annotations (**Figure 2C**) and 71.6% of proteins match their localization annotation in thapsigargin-treated cells **(Supplemental Figure S5**). From the associated SILAC data of the proteins with spatial information, we further quantified and compared the turnover kinetics of 2516 proteins (**Supplemental Data S2**); hence we were able to acquire proteome-wide spatial and temporal information in matching samples from a single experiment.

**Figure 2.**
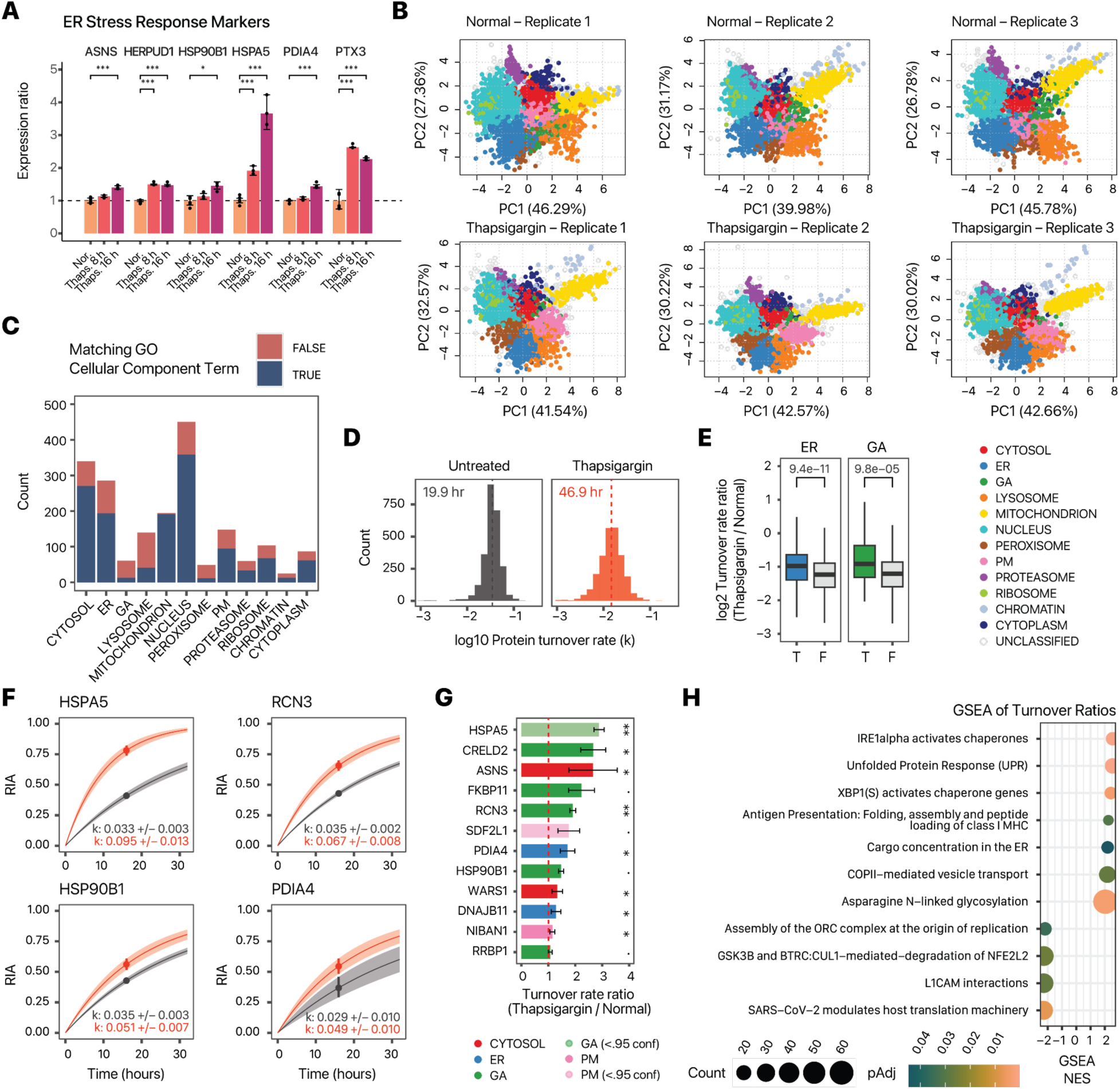
Simultaneous measurements of spatial and temporal kinetics under UPR. **A.** Bar charts showing activation of known ER stress markers upon thapsigargin treatment for 8 hours and 16 hours. X-axis: ER stress markers; y-axis: expression ratio (n=6 normal AC16; n=3 thapsigargin). *: limma adjusted P < 0.01; **: limma adjusted P < 0.001; ***: limma adjusted < 0.0001; error bars: s.d. **B.** PC1 and PC2 of proteins spatial map showing the localization of confidently allocated proteins in normal and thapsigargin-treated AC16 cells. Each data point represents a protein; color represents classification of subcellular localization. **C.** Distribution of light (unlabeled) protein features in each of the 12 subcellular compartments (n=3); fill color represents whether the protein is also annotated to the same subcellular compartment in UniProt Gene Ontology Cellular Component terms. **D.** Histograms of the determined log10 protein turnover rates in control and thapsigargin treated cells (n=3). Text overlay indicates median half-life. **E.** Boxplot showing the log2 turnover rate ratios in thapsigargin over normal AC16 cells for proteins that are localized to the ER (blue) (T) or not (F); or the Golgi (GA; green). P values: Mann-Whitney test. A Bonferroni corrected threshold of 0.05/13 is considered significant. Center line: median; box limits: interquartile range; whiskers: 1.5x interquartile range. **F.** Example of best fit curves in the first-order kinetic model at the protein level between normal (gray), and thapsigargin treated (red) AC16 cells showing four known ER stress markers with elevated turnover (HSPA5, RCN3, HSP90B1, and PDIA4). Because the sampling time point is known, the measured relative isotope abundance of a peptide (prior to reaching the asymptote) is sufficient to define the kinetic curve and the parameter of interest (k). **G.** Turnover rate ratio (thapsigargin vs. normal) of the top proteins with elevated temporal kinetics in UPR within the ER (blue) and Golgi (green); .: Mann-Whitney test FDR adjusted P value < 0.1; *: < 0.05; ** < 0.01; red dashed line: 1:1 ratio; bars: standard error. **H.** Gene set enrichment analysis (GSEA) of turnover rate ratios in thapsigargin treatment; proteins with faster kinetics are significantly enriched in DNA damage response and UPR pathways. Color: FDR adjusted P values in GSEA; x-axis: GSEA normalized enrichment score (NES). Size: number of proteins in gene set.

Considering the temporal kinetics data, we observed a proteome-wide decrease in fractional synthesis rates under thapsigargin challenge compared with normal cells (median protein half-life 46.9 vs. 19.9 hours; Mann-Whitney test P < 2.2e–16) (**Figure 2D**). This slowdown is consistent with the extensive shutdown in protein translation due to ribosome remodeling under integrated stress response (Bresson et al., 2020; Pakos-Zebrucka et al., 2016), shown here by the decreased rate of SILAC incorporation into proteins. Notwithstanding the overall slowdown, we also observed a wide range of protein turnover rates in both the untreated and thapsigargin treated conditions that differ by the assigned subcellular compartment (**Supplemental Figure S6**). Changes in protein kinetics following thapsigargin also varies by compartment, with ER and Golgi proteins having significantly less slowdown of protein kinetics compared to protein in other compartments (Mann-Whitney test P: 9.4e–11 and 9.8e–5, respectively; << 0.05/13) (**Figure 2E**). On an individual protein level, out of the 2516 proteins measured, 1542 showed significant changes in temporal kinetics (Mann-Whitney test, FDR adjusted P value < 0.1), but the vast majority of these proteins show decreases in turnover as expected, with only 12 proteins showing significant increased temporal kinetics. Among these are the induced ER stress markers BiP/HSPA5, HSP90B1, and PDIA4 (**Figure 2F; Supplemental Data S3**) but also other ER and Golgi proteins that may be involved in stress response (**Figure 2G**). SDF2L1 (stromal-cell derived factor 2 like 1) is recently described to form a complex with the ER chaperone DNAJB11 to retain it in the ER (Hanafusa et al., 2019). In control cells, we found that SDF2L1 has a basal turnover rate of 0.027/hr. Upon thapsigargin treatment, its turnover rate increased to 0.048 /hr (adjusted P: 0.07). DNAJB11 also experienced accelerated kinetics (1.28-fold in thapsigargin, adjusted P 0.029) hence both proteins may be preferentially synthesized during UPR. On a proteome level, gene set enrichment analysis (GSEA) of temporal kinetics changes show a preferential enrichment of proteins in unfolded protein response (FDR adjusted P: 4.1e–4), ER to Golgi anterograde transport (FDR adjusted P 0.036) and N-linked glycosylation (FDR adjusted P: 1.7e–3) but a negative enrichment of translation-related terms (**Figure 2H**). Overall, protein kinetic changes are modestly correlated with protein abundance changes (**Supplemental Figure S7**), suggesting that AC16 cells actively regulate protein synthesis and degradation kinetics in normal and stressed conditions beyond changes in protein abundance.

### Changes in protein subcellular distribution under ER stress

We next analyzed the spatial proteomics component of the data to find proteins that change in their subcellular localization following thapsigargin treatment. To do so, we used a Bayesian statistical model implemented in the BANDLE package to estimate the differential spatial localization of proteins. In total, we identified 1,306 translocating protein features (687 light and 619 heavy) in thapsigargin under a stringent filter of BANDLE differential localization probability > 0.95 with an estimated FDR of 0.0018 (0.18%), and further filtered using a bootstrap differential localization probability of > 0.95. We then further prioritized 330 pairs of differentially localized proteins where the light and heavy features both show confident differential localization (**Supplemental Data S2**, **Supplemental Data S4**). The differential localization of these 330 proteins recapitulate previously established relocalization events in cellular stress response, capturing the migration of caveolae toward the mitochondrion under cellular stress (Fridolfsson et al., 2012) (**Supplemental Figure S8A)**, and the engagement of EIF3 to ribosomes in EIF3-dependent translation initiation in integrated stress response (Guan et al., 2017) (**Supplemental Figure S8B**), thus supporting the confidence of the spatial translocation assignment.

From the results, we discerned three major categories of differential localization behaviors in ER stress that revealed new insights into proteome-wide features of UPR. First, we observed the externalization of proteins toward the plasma membrane (**Figure 3A**). The large neutral amino acid transporter component SLC3A2 is localized to the lysosome fraction in normal cells (Pr > 0.999) but in thapsigargin-treated cells is localized confidently to the plasma membrane (BANDLE differential localization probability > 0.999) (**Figure 3B**). Showing similar behaviors are SLC7A5, the complex interacting partner of SLC3A2; and two other amino acid transporters SLC1A4, and SLC1A5 (**Figure 3C**); whereas the ion channel proteins SLC30A1, ATP1B1, ATB1B3 and ATP2B1 also showed confident localization toward the cell surface (**Figure 3C**). The change in localization of SLC3A2 is corroborated by immunostaining (**Figure 3D**), which shows a decrease in co-localization between immunostaining signals of SLC3A2 and lysosome marker LAMP2 upon thapsigargin treatment (**Figure 3E**).

**Figure 3.**
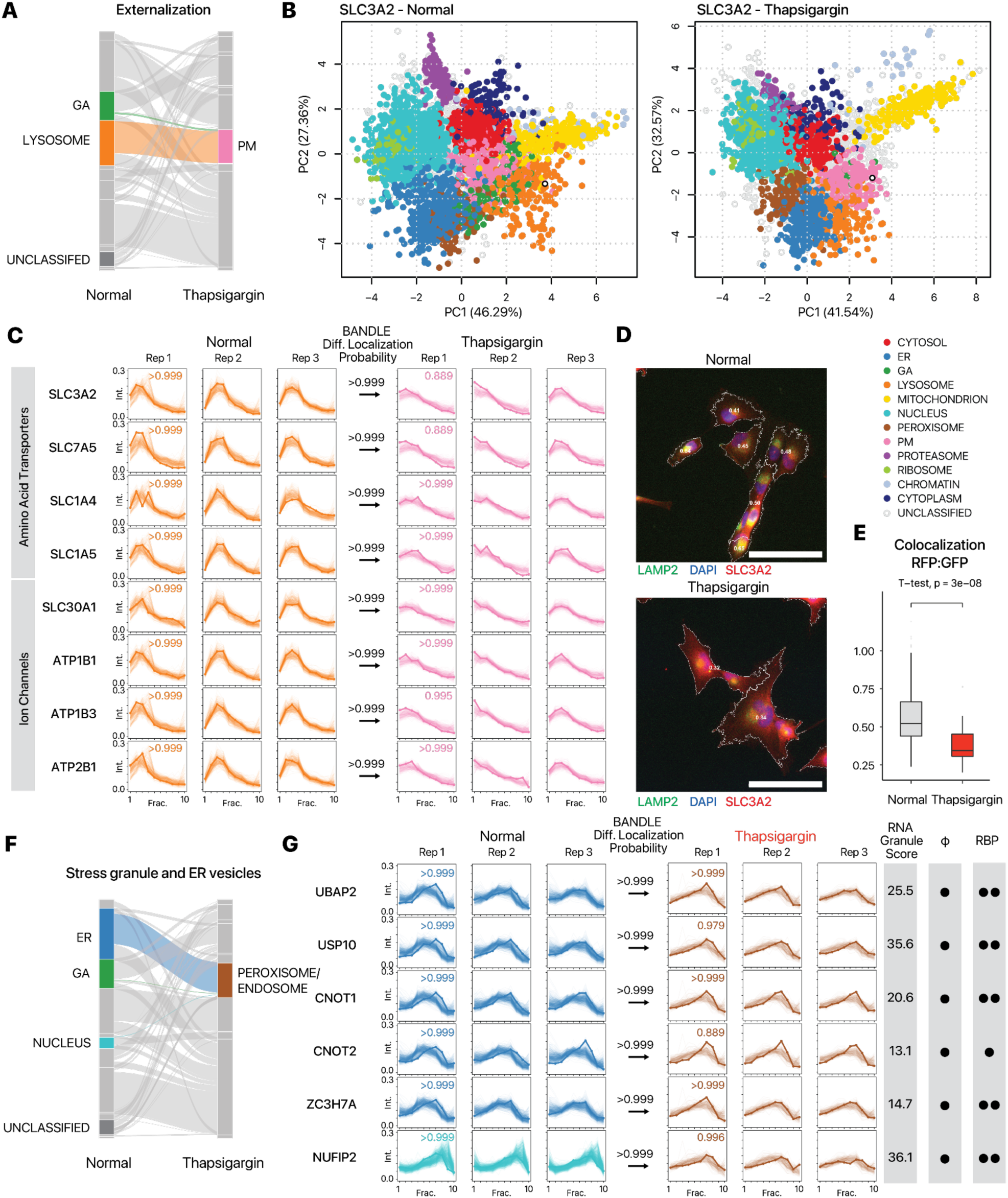
SPLAT captures extensive protein translocation in AC16 cells under UPR. **A.** Alluvial plot of translocation events (> 0.99 BANDLE translocation probability; estimated FDR < 1%) following thapsigargin treatment showing a cohort of proteins moving from the Golgi apparatus (GA) and lysosome towards the plasma membrane (PM) (n=3). **B.** Protein spatial map for SLC3A2 (open black circle) in normal (left) and thapsigargin-treated (right) AC16 cells, showing its colocalization with lysosomal proteins in normal cells and in PM proteins in thapsigargin-treated cells. Colors represent allocated subcellular localization. **C.** Ultracentrifugation fraction profile of SLC3A2 and other amino acid transporters SLC7A5, SLC1A4, SLC1A5 and ion channel proteins SLC30A1, ATP1B1, ATP1B3, and ATP2B1 with similar migration patterns. X-axis: fraction 1 to 10 of ultracentrifugation. Y-axis: relative channel abundance. Bold lines represent the protein in question; light lines represent ultracentrifugation profiles of all proteins classified to a respective localization. Colors correspond to subcellular localization in panel B and for all AC16 data throughout the manuscript; numbers within boxes correspond to BANDLE allocation probability to compartment. **D.** Immunofluorescence of SLC3A2 (red) against the lysosome marker LAMP2 (green) and DAPI (blue). Numbers in cell boundary: colocalization score per cell. Scale bar: 90 µm. **E.** Colocalization score (Mander’s correlation coefficient) between SLC3A2 and LAMP2 decreases significantly (two-tailed unpaired t-test P value: 3e–8) following thapsigargin treatment, consistent with movement away from lysosomal fraction (n= 205 normal cells, n = 32 thapsigargin treated cells). Center line: median; box limits: interquartile range; whiskers: 1.5x interquartile range; points: outliers. **F.** Alluvial plot showing the migration of ER, GA, and nucleus proteins toward the peroxisome/endosome containing fraction in thapsigargin treated cells. **G.** Ultracentrifugation fraction profile of stress granule proteins UBAP2, USP10, CNOT1, CNOT2, ZC3H7A, and NUFIP2. RNA Granule Score: score from RNA Granule Database (https://rnagranuledb.lunenfeld.ca/). A score of 7 or above is considered a known stress granule protein. Phi: predicted phase separation participation. Circle denotes a prediction of True within the database. RBP: Annotated RNA binding protein on the RNA Granule Database. One circle denotes known RBP in at least one data set; two circles denote known RBP in multiple datasets.

Second, thapsigargin treated AC16 cells are associated with an increase in proteins classified into the peroxisome fraction including proteins whose locations changed from the ER, Golgi, and the nucleus in normal cells (**Figure 3F**). In mammalian cells, ER and peroxisomes are spatially adjacent; the peroxisome associated fractions sediment prominently at 5000–9000 × g (F3 and F4) in the LOPIT-DC protocol (**Supplemental Figure S2**), marked by canonical peroxisome markers PEX14 and ACOX1 (**Supplemental Data S2**). However, although this compartment was trained using peroxisome markers, the majority of proteins categorized into this compartment are not annotated to be in the peroxisome whereas 30 out of 49 (61%) of proteins in the control cells allocated to this compartment were annotated also as endosome, including canonical markers EEA1 and VPS35L. We thus refer to this compartment hereafter as peroxisome/endosome. Moreover, proteins that become differentially localized to this fraction in thapsigargin include known stress granule proteins UBAP2, USP10, CNOT1, CNOT2, CNOT3, CNOT7, CNOT10, ZC3H7A, and NUFIP2 (**Figure 3G and Supplemental Figure S9)**, which show high-confidence translocation to the peroxisome/endosome fraction, and are known RNA binding proteins that participate in phase separation, consistent with stress granule formation in UPR. Notably, LMAN1, LMAN2, SCYL2, and SNX1 are RNA-binding proteins that are not currently established stress granule components and show identical translocation patterns, nominating them as potential participants in RNA granule related processes in AC16 cells for further studies (**Supplemental Figure S9**). Other proteins in this fraction include the ER-to-Golgi transport vesicle proteins GOLT1B, GOSR2, RER1, and NAPA (**Supplemental Figure S9**).

Lastly, we see evidence of proteins from the ER and Golgi targeted to the lysosome (see Tunicamycin section below). Thus taken together, the spatial proteomics component of the data reveals a complex network of changes in protein spatial distribution during UPR.

### Partition of newly synthesized and pre-existing protein pools

We next considered the interconnectivity of temporal and spatial dynamics, namely whether some localization changes are contingent upon protein pool lifetime, such as where light (old) protein does not change in spatial distribution but the heavy (new) proteins display differential translocation upon UPR. Because the spatial profiles of the light and heavy proteins are acquired independently, this experimental design allowed us to examine whether old and new proteins are localized to identical cellular locales. In both normal and thapsigargin-treated cells, we found that the independently measured spatial profiles of light (pre-existing) proteins and their corresponding heavy SILAC (newly synthesized) counterparts are highly concordant, with a normalized spatial distribution distance (see Supplemental Methods) of 0.020 [0.015 – 0.030], compared to 0.117 [0.080 – 0.155] in random pairs of pre-existing proteins (1,614 light-heavy pairs, Mann-Whitney P < 2.2e–16) in normal cells, and 0.028 [0.019 – 0.041] and 0.122 [0.081 – 0.161] in thapsigargin-treated cells (1,614 light-heavy pairs, Mann-Whitney P < 2.2e–16) (**Figure 4A**). This robust agreement provided an additional independent confirmation on the accuracy of the spatial measurements. Consistently, among heavy-light protein pairs with confidently assigned subcellular localization, the heavy and light proteins are assigned to the identical subcellular compartment in 93% and 89% of the cases in normal and thapsigargin-treated AC16 cells, respectively (**Figure 4B**). We focused on the unusual cases where the spatial distribution distances between the heavy and light proteins increased noticeably following thapsigargin treatment, as they may be indicative of localization changes that are dependent upon time since synthesis. These include two proteins EGFR and ITGAV with an uncommon increase in heavy-light spatial distances (Z: 12.00 and 2.24, respectively) (**Figure 4C**). Epidermal growth factor receptor (EGFR/ErbB1/HER1) is a receptor tyrosine kinase with multiple subcellular localizations and signaling roles, and is implicated in cardiomyocyte survival (Lee et al., 2020). Following a variety of stressors, EGFR is known to be inactivated by intracellular trafficking, including being internalized to the early endosome and lysosome following oxidative stress and hypoxia in cancer cells (Tan et al., 2016). In the spatial proteomics data, the spatial distribution of EGFR borders the lysosome and plasma membrane fractions, which we interpret as EGFR having potential multiple pools including a cell surface fraction (**Figure 4D**). In the thapsigargin treated cells, the light-heavy spatial distance of EGFR increased from 0.014 in normal cells to 0.052, and the dynamic SILAC labeled pool (heavy) becomes internalized toward the ER (BANDLE differential localization probability: >0.999) but not the pre-existing (light) pool (**Figure 4E**). The data therefore suggests that the internalization of EGFR under thapsigargin is likely to be due to endomembrane stalling or redistribution upon new protein synthesis, possibly leading to fewer new EGFR molecules reaching the cell surface. Likewise, in thapsigargin-treated AC16 cells, newly-synthesized ITGAV (integrin subunit alpha V) shows a partition from the plasma membrane fraction to the ER fraction but not the old/existing protein pool, concomitant with an increase in spatial distribution distance from 0.017 to 0.057 between old and new proteins (**Figure 4F–G**). With the function of integrins as cell surface receptors that function in intracellular-to-extracellular and retrograde communication, the ER localization of newly synthesized ITGAV, such as due to stress-induced stalling of protein trafficking along the secretory pathway, could indicate a decrease in integrin signaling function through spatial regulation rather than protein abundance. To partially corroborate the partial redistribution of EGFR, we performed immunocytochemistry imaging of EGFR subcellular distribution in AC16 cells with or without thapsigargin (**Figure 4H**). Thapsigargin treatment did not increase cell size (**Figure 4I**), and whereas there is an increase in immunofluorescence signal of EGFR in thapsigargin (**Figure 4J**), this signal is distributed preferentially to the interior of the cell such that there is a significant reduction in the ratio of mean intensity at cell borders over the whole cell in thapsigargin vs. untreated cells (0.673 vs. 0.746, n=93 and 71 cells, Mann-Whitney P: 1.7e–4) (**Figure 4K**), consistent with a partial redistribution of EGFR toward an internal pool. Taken together, these examples demonstrate the SPLAT strategy can be used to distinguish time-dependent differential localization of proteins such as due to the trafficking of newly synthesized proteins.

**Figure 4.**
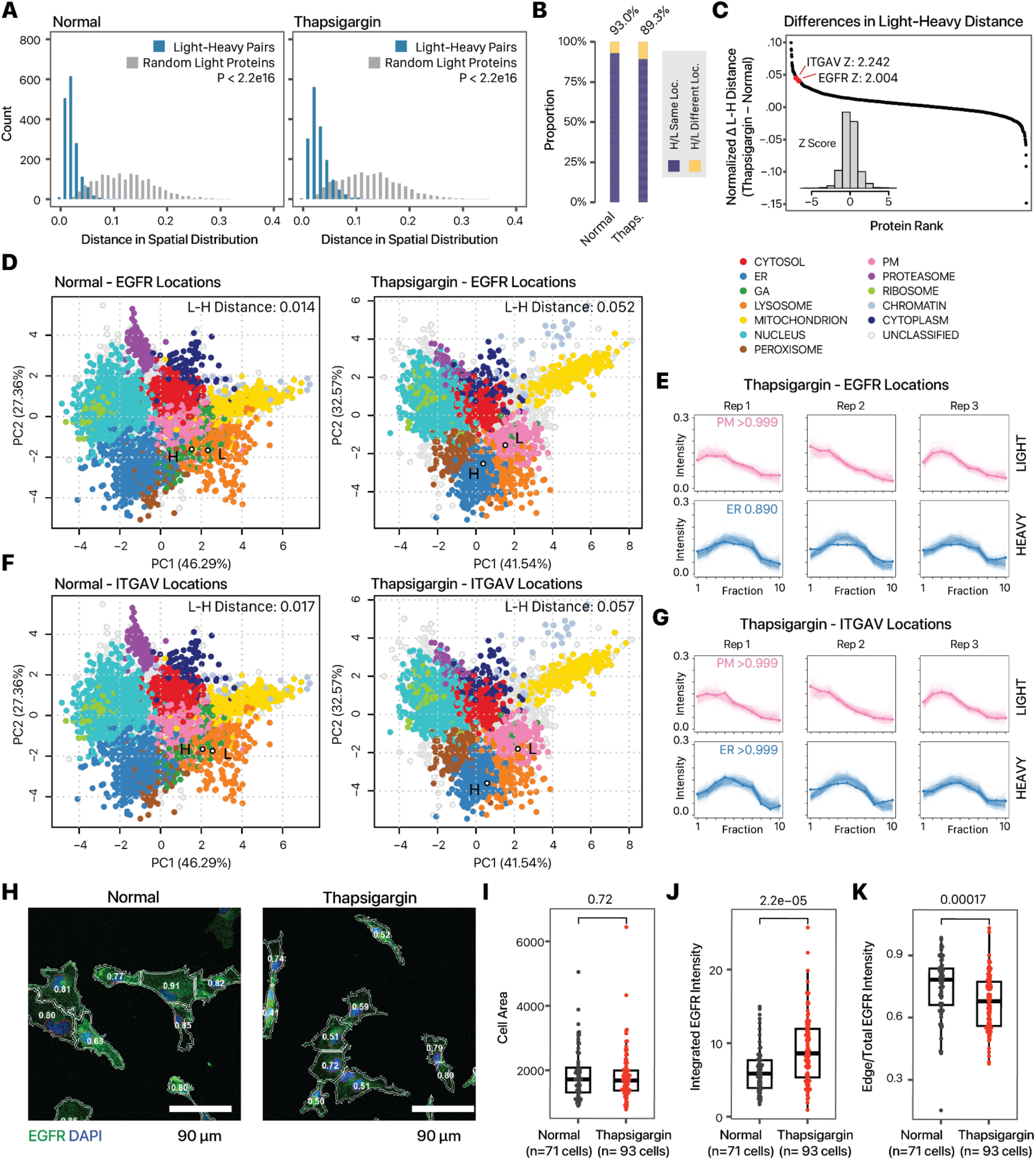
SPLAT reveals protein-lifetime dependent translocation. **A.** Histogram showing the similarity in light and heavy proteins in normalized fraction abundance profiles in (left) normal and (right) thapsigargin-treated AC16 cells. X-axis: the spatial distribution distance of two proteins is measured as the average euclidean distance of all TMT channel relative abundance in the ultracentrifugation fraction profiles across 3 replicates; y-axis: count. Blue: distance for 1,614 quantified light-heavy protein pairs (e.g., unlabeled EGFR, heavy SILAC-labeled EGFR). Grey: distribution of each corresponding light protein with another random light protein. P value: Mann-Whitney test. **B.** Proportion of heavy-light protein pairs with confidently assigned localization that are assigned to the same location (purple) in normal (left; 93%) and thapsigargin-treated (right; 89%) cells. **C.** Ranked changes in heavy-light pair euclidean distance upon thapsigargin treatment. The difference in heavy-light distances in thapsigargin is adjusted by the average changes in the spatial distance of the light protein with 250 other sampled light proteins to calculate the normalized difference. The majority of proteins show no change (+/- 0.02 in euclidean distance). The positions of EGFR and ITGAV are highlighted. Inset: Z score distribution of all changes. **D.** Spatial map showing the location of the light and heavy EGFR in normal and thapsigargin-treated AC16 cells. Each data point is a light or heavy protein species. Colors correspond to other AC16 experiments in the manuscript. Numbers correspond to euclidean distance in fraction profiles over 3 replicates. **E.** Corresponding fraction profiles; x-axis: ultracentrifugation fraction; y-axis: fractional abundance. Post-labeling synthesized EGFR is differentially distributed in thapsigargin and shows ER retention (blue), whereas the preexisting EGFR pool remains to show a likely cell surface localization (pink) after thapsigargin. **F-G**. As above, for ITGAV. **H.** Confocal imaging of EGFR immunofluorescence supports a partial relocalization of EGFR from the cell surface toward internal membranes following thapsigargin treatment. Numbers: The mean intensity of the labeled EGFR channel of a 3 pixel border at cell boundaries was divided by mean intensity of the whole cell to estimate the ratio of EGFR at the plasma membrane to the cell interior. Blue: DAPI; Green: EGFR; scale bar: 90 µm. **I.** Cell areas; Mann-Whitney P: 0.72. Center line: median; box limits: interquartile range; whiskers: 1.5x interquartile range. **J.** Total EGFR intensity per cell; Mann-Whitney P: 2.2e-05. Center line: median; box limits: interquartile range; whiskers: 1.5x interquartile range. **K.** Edge/total intensity ratios in normal and thapsigargin-treated AC16 cells (n=71 normal cells; n=93 thapsigargin cells; Mann-Whitney P: 1.7e–4). Center line: median; box limits: interquartile range; whiskers: 1.5x interquartile range.

### Spatiotemporal proteomics highlights similarities and differences of ER stress induction protocols

We next investigated the protein spatiotemporal features of AC16 cells under the treatment of tunicamycin, another compound commonly used to induce ER stress in cardiac cells (Liu et al., 2012; Toro et al., 2022) by inducing proteostatic stress via inhibition of nascent protein glycosylation. Three biological replicate SPLAT experiments were performed in tunicamycin-treated cells to resolve the temporal kinetics and subcellular localisation of proteins (**Supplemental Figures S1, S2C, S5B, S6, S7B, S10**). Tunicamycin treatment at 1 µg/mL for 16 hours robustly induced the known ER stress response markers BiP/HSPA5, HSP90B1, PDIA4, CALR, CANX, and DNAJB11 (limma FDR adjusted P < 0.10) (**Figure 5A**) demonstrating effective ER stress induction. Overall, tunicamycin treatment led to a lesser slowdown of temporal kinetics than thapsigargin (average protein half-life 32.6 hours) (**Figure 5B; Supplemental Data S5**). As in thapsigargin treatment, the kinetic changes following tunicamycin are different across cellular compartments, with ER and Golgi proteins having relatively faster kinetics than other cellular compartments (Mann-Whitney test P: 1.3e–13 and 6.6e–5; << Bonferroni corrected threshold 0.05/13); whereas the greatest reduction was observed among proteins localized to the lysosome, a compartment closely linked to glycosylation and recycling of glycans (Mann-Whitney test P: 1.8e–10) (**Figure 5C**). Gene set enrichment analysis (GSEA) of turnover rate ratios revealed a significant positive enrichment of UPR proteins (adjusted P 3.5e–3) and DNA repair terms (e.g., processing of DNA double-strand break ends; adjusted P 0.017) and a negative enrichment of translation related terms (**Figure 5D**). Compared to thapsigargin treatment however, no significant enrichment of glycosylation and vesicle transport related terms were found in tunicamycin. Inspection of individual protein kinetics changes likewise revealed both similar induction of the ER stress response markers HSPA5, HSP90B1 and PDIA4 as in thapsigargin treatment, but other stress response genes PDIA3 and NIBAN1 are not induced in thapsigargin (**Figure 5E**). On the other hand, RCN3 (reticulocalbin 3) is an ER lumen calcium binding protein that regulates collagen production (Martínez-Martínez et al., 2017) and shows increased temporal kinetics in thapsigargin (**Figure 2F**) but not in tunicamycin (ratio 0.76 over normal; **Supplemental Data S5**), altogether reflecting potential differences in stress response modality to a different ER stress inducer.

**Figure 5.**
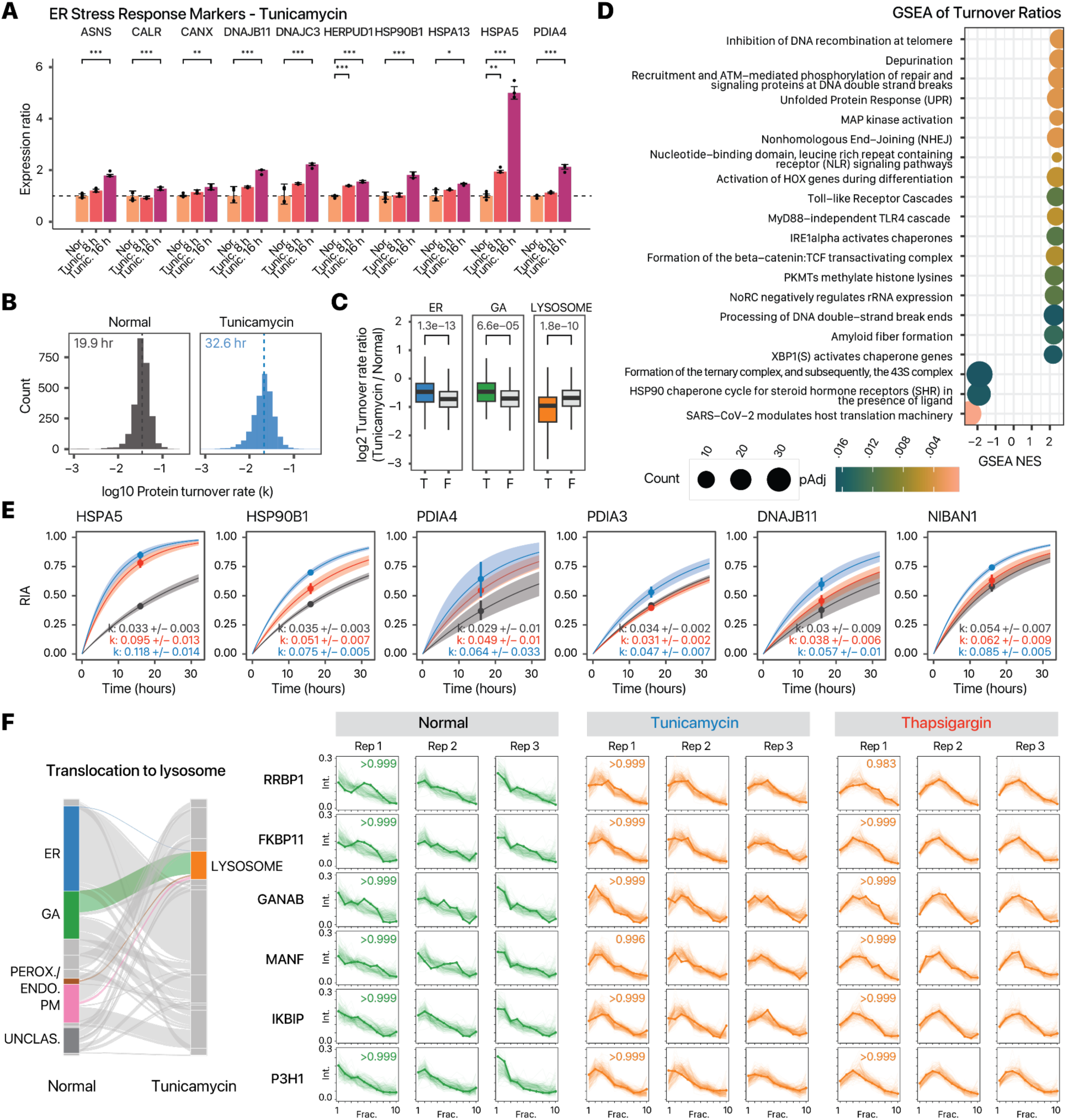
Comparison of ER stress induction methods. **A.** Bar charts showing activation of known ER stress markers upon tunicamycin treatment for 8 hours and 16 hours. X-axis: ER stress markers; y-axis: expression ratio (n=6 normal AC16; n=3 tunicamycin). *: limma adjusted P < 0.01; **: limma adjusted P < 0.001; ***: limma adjusted P < 0.0001; error bars: s.d.. **B.** Histograms of the determined log10 protein turnover rates in control and tunicamycin treated cells (n=3). **C.** Boxplot showing the log2 turnover rate ratios in tunicamycin over normal AC16 cells for proteins that are localized to the ER (T) or not (F); Golgi apparatus, or the lysosome. P values: two-tailed t-test. Center line: median; box limits: interquartile range; whiskers: 1.5x interquartile range. A Bonferroni corrected threshold of 0.05/13 is considered significant. **D.** Gene set enrichment analysis (GSEA) of turnover rate ratios in tunicamycin treatment. Color: FDR adjusted P values in GSEA; x-axis: GSEA normalized enrichment score (NES). Size: number of proteins in the gene set. **E.** Example of best fit curves in the first-order kinetic model at the protein level between normal (gray), tunicamycin (blue) and thapsigargin (red) treated AC16 cells showing known ER stress markers with elevated turnover in both ER stress inducers (HSPA5, HSP90B1, and PDIA4) as well as stress response proteins with elevated turnover only in tunicamycin (PDIA3, DNAJB11, NIBAN1). **F.** Alluvial plot showing the migration of ER, GA, and peroxisome/endosome proteins toward the lysosome (left). On the right, the ultracentrifugation fraction profiles of translocating proteins RRBP1, FKBP11, GANAB, MANF, IKBIP, and P3H1 are shown that are targeted toward the lysosome in both tunicamycin and thapsigargin treatment (BANDLE differential localization probability > 0.95). Numbers in boxes are the BANDLE allocation probability in each condition (n=3).

Parallel to the less prominent changes in vesicle transport, tunicamycin treatment also led to fewer translocating proteins than thapsigargin, with 620 translocating features (including 282 light proteins and 338 heavy proteins) at BANDLE differential localization probability > 0.95, corresponding to an estimated FDR of 0.35%, and thresholded by bootstrapping differential localization probability > 0.95; from which we highlighted 109 proteins where the heavy and light versions both showed translocation. The spatial data revealed a high degree of similarity but also notable differences with thapsigargin-induced ER stress. We found that in both tunicamycin and thapsigargin treatment, there was evidence of lysosome targeting from other endomembrane compartments, including: RRBP1, a ribosome-binding protein of the ER, GANAB, a glucosidase II alpha subunit integral to the proper folding of proteins in the ER, FKBP11, a peptidyl-prolyl cis/trans isomerase important to the folding of proline-containing peptides, IKBIP, an interacting protein to the IKBKB nuclear kinase, and MANF, a neurotrophic factor which has relations to ER stress-related cell death when its expression is lowered (BANDLE differential localization probability > 0.999) (Sayers et al., 2022) (**Figure 5F**). Among these proteins was the collagen synthesis enzyme P3H1 in both thapsigargin and tunicamycin. Interestingly, prior work found no correlation between the protein abundance of collagen modifying enzymes with the known reduction of collagen synthesis in chondrocytes and fibroblasts under ER stress (Vonk et al., 2010). The results here suggest that the functional decline may instead correlate with a change in the subcellular localization of collagen modifying enzymes in AC16 cells. Tunicamycin treatment also induced old-new protein partitions in EGFR and ITGAV as observed in thapsigargin (**Supplemental Figure S11**).Notably, although tunicamycin also induced the translocation of proteins toward the peroxisome/endosome fraction, different proteins are involved, including the stress response proteins DNAJB11, DNAJC3, DNAJC10, and PDIA6 as well as other proteins EMC4, EMC8, VAPA, and VAPB (**Supplemental Figure S12**) which further outlines different modalities of cellular response toward two different ER stress inducers. The translocating stress response proteins DNAJB11, DNAJC10, and PDIA6 also showed significant acceleration in temporal kinetics in tunicamycin (Mann-Whitney test, FDR adjusted P < 0.10; **Supplemental Data S6**) which is consistent with specific production of the proteins followed by shuttling to subcellular location for their function during stress response.

### Application of SPLAT to the mechanism of cardiotoxicity in iPSC-CM models

We next assessed the applicability of SPLAT toward a different, non-proliferating cell type (human iPSC-derived cardiomyocytes [iPSC-CMs]) and its utility for interrogating the mechanism of cardiotoxicity following proteasome inhibitor treatment (**Figure 6A**). The ubiquitin proteasome system is responsible for the degradation of over 70% of cellular proteins. Compounds that inhibit proteasome function, including carfilzomib, are widely used as cancer treatment and have led to remarkable improvement in the survival of multiple myeloma patients. Mechanistically, carfilzomib functions by binding to and irreversibly inhibiting the proteasome catalytic subunit PSMB5 (Cromm and Crews, 2017), leading to the accumulation of unfolded proteins in cancer cells. Importantly, despite its efficacy as a cancer treatment, carfilzomib also leads to cardiotoxic adverse effects including heart failure (<20%), arrhythmia (<10%), and hypertension (11-37%) in a significant number of patients (IBM Watson Health, 2023). This cardiotoxicity has been modeled in vitro by exposure of 0.01 – 10 µM carfilzomib to human iPSC-CMs (Forghani et al., 2021), yet the molecular mechanisms of carfilzomib cardiotoxicity remain incompletely understood. To examine the protein spatiotemporal changes upon carfilzomib-mediated proteasome inhibition in cardiac cells, we differentiated contractile iPSC-CMs using a small molecule based protocol, and treated the cells with 0.5 µM carfilzomib. To verify toxicity modeling, we measured iPSC-CM viability and phenotypes under 0 to 5 µM carfilzomib for 24 and 48 hours. Under the chosen treatment (0.5 µM for 48 hours), iPSC-CMs showed sarcomeric disarray consistent with prior observations on the cardiotoxic effects of carfilzomib (**Figure 6B**) but maintained viability (82%) (**Figure 6C**), while showing significant decreases in oxygen consumption (**Figure 6D**), basal respiration (**Figure 6E)**, and maximal respiration (**Figure 6F**), whereas higher doses are accompanied with drops in viability at 48 hours and an increase in proton leak (**Figure 6G**). ATP production at the 0.5 µg dose was significantly lower than untreated cells at both 24 and 48 hours (**Figure 6H**). Therefore cardiotoxicity due to carfilzomib can be recapitulated in a human iPSC-CM model, consistent with prior work in the literature.

**Figure 6.**
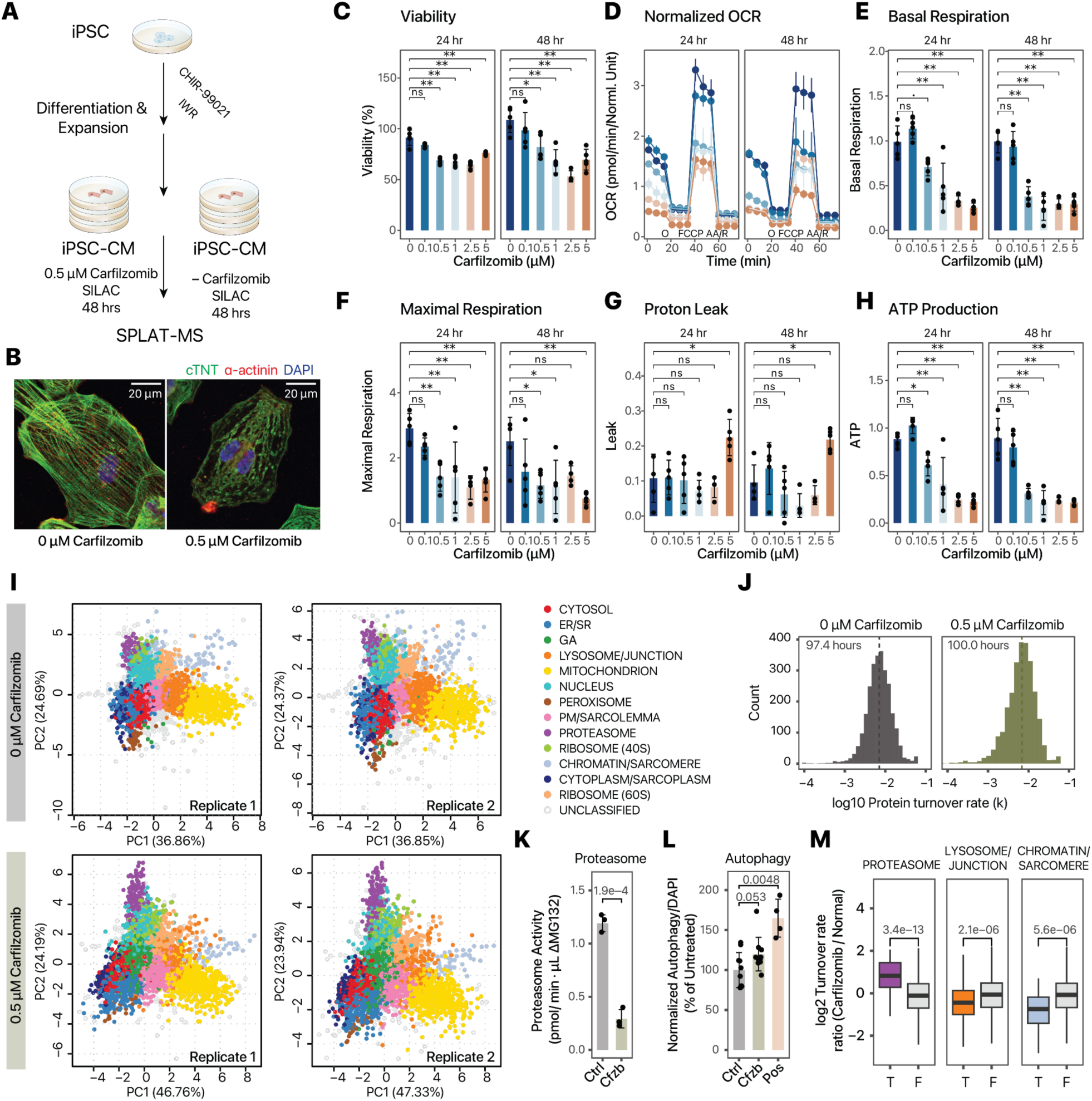
Applicability in human iPSC-derived cardiomyocytes. A. Schematic of human iPSC differentiation into cardiomyocytes, carfilzomib treatment, and SPLAT analysis. **B.** Confocal microscopy images showing sarcomeric disarray in iPSC-CMs upon 48 hrs of 0.5 µM carfilzomib; green: cTNT, red: alpha-actinin; blue: DAPI; scale bar: 20 µm. **C–H.** Cell viability (%), normalized Seahorse oxygen consumption rate (OCR; pmol/min), basal respiration, maximal respiration, proton leak, and ATP production upon 0 – 5 µM carfilzomib for 24 or 48 hrs; .: adjusted P < 0.1; *: adjusted P < 0.05; **: adjusted P < 0.01, ANOVA with Tukey’s HSD post-hoc at 95% confidence level; n=5. Error bars: s.d. for bar charts in panels C, E, F, G, H; s.em. for the OCR graph in panel D. Colors in panel D: dosage, same as panel C. O: Oligomycin; AA/R: Antimycin A/Rotenone. **I.** Spatial map with 13 assigned subcellular localizations in iPSC-CMs at the baseline (top) and upon 0.5 µM carfilzomib treatment (n=2). **J.** Histogram of log10 protein turnover rates (k), with median half-life of 97.4 hours and 100.0 hours in normal and carfilzomib-treated iPSC-CM. **K.** Proteasome activity in iPSC-CMs treated with 0 (Ctrl) vs. 0.5 µM carfilzomib (Cfzb) for 48 hrs. P value: two-tailed t-test; n = 3. **L.** Autophagy assay for iPSC-CMs treated with 0 (Ctrl) vs. 0.5 µM carfilzomib (Cfzb) for 48 hrs, and positive control (Pos); data were normalized to DAPI and normal cells. P value: two-tailed t-test; n = 10. **M.** log2 Turnover rate ratios between carfilzomib-treated and untreated iPSC-CM from the spatiotemporal proteomics data. Proteins assigned the proteasome compartment have significantly increased temporal kinetics; proteins in the lysosome/junction and chromatin/sarcomere compartments have significantly reduced temporal kinetics. P values: Mann-Whitney; with a threshold of 0.05/14 considered significant. Center line: median; box limits: interquartile range; whiskers: 1.5x interquartile range.

From the untreated and carfilzomib-treated (0.5 µM, 48 hrs) human iPSC-CMs, we constructed a protein subcellular spatial map that takes into account several features of the iPSC-CM cell type, including the inclusion of cell junction and desmosome proteins, as well as a sarcomere protein compartment, that are not apparently recognized as discrete compartments in the prior spatial maps (**Supplemental Figure S13**). In addition, the 40S and 60S ribosomes showed clear separation in iPSC-CM unlike in AC16 cells and are hence classified separately. This separation is consistent with less active protein translation in this cell type. In total, we mapped the subcellular localization of 5,047 protein features including 2,680 old proteins and 2,367 new proteins using a stringent two-peptide filter at 1% FDR of protein identification, including 2,010 old proteins and 1,737 new proteins assigned to one of 13 subcellular localization with >95% confidence after removing outliers (**Figure 6I; Supplemental Data S7**). The iPSC-CM spatial map achieved similar levels of concordance with known cellular component annotations as in AC16 cells (70.8% with known annotation matching the assigned compartment in normal iPSC-CM; 63.0% in carfilzomib-treated cells) (**Supplemental Figure S14**). The iPSC-CM map has comparable spatial distance between light and heavy protein pairs as in AC16 cells, and 87.6% heavy-light protein pairs map to the same compartment in the baseline, supporting that there is sufficient spatial resolution to resolve subcellular compartment differences in this cell type (**Supplemental Figure S14**).

Among proteins with spatial information, we compared the temporal kinetics of 2,648 proteins. Unexpectedly, there was no overall slowdown of protein temporal kinetics with the median protein half-lives being 97.4 hours and 100.0 hours in normal and carfilzomib-treated cells, respectively (**Figure 6J**; **Supplemental Data S8**), suggesting that at 48 hours following proteasome inhibitor treatment, the observed cellular toxicity is not directly explainable by a drop of per-protein average in global protein degradation. At 48 hours of carfilzomib treatment in iPSC-CMs, proteasome chymotrypsin-like activities are partially suppressed but significant partial activities are also observable (**Figure 6K**); whereas other proteolysis mechanisms may also compensate for proteasome inhibition, including a suggestive increase in autophagy (P: 0.053) (**Figure 6L**). Inspection of the spatial data revealed that the changes in protein kinetics upon carfilzomib are localization specific, with a significant reduction in chromatin/sarcomere protein turnover rate, and significant increase for the proteasome compartment under carfilzomib treatment (**Figure 6M**).

Notably, on an individual protein level we find that the majority of proteins with increased protein kinetics belong to subunits of the regulatory 19S complex rather than the core 20S complex (**Figure 7A**) suggesting possible changes in 26S proteasome activity and target engagement. In addition to proteasome subunits, the temporal kinetics revealed a robust induction of chaperons HSP90AA1/HSP90A, HSP90AB1/HSP90B, HSPA4, and BAG3; and ERAD associated proteins VCP and UFD1 (**Figure 7B, Supplemental Data S8**). Within the mitochondrion, quality control proteins HSPD1, HSPE1, and CLPB are induced (**Figure 7A**). In contrast, among proteins that show reduced turnover in carfilzomib treatment are major sarcomeric proteins MYH6, MYH7, MYBPC3, MYL7; as well as proteins classified to the cell junction compartment dystrophin (DMD) and utrophin (UTRN), and the desmosome complex protein desmoplakin (DSP) (**Figure 7A & C**).

**Figure 7.**
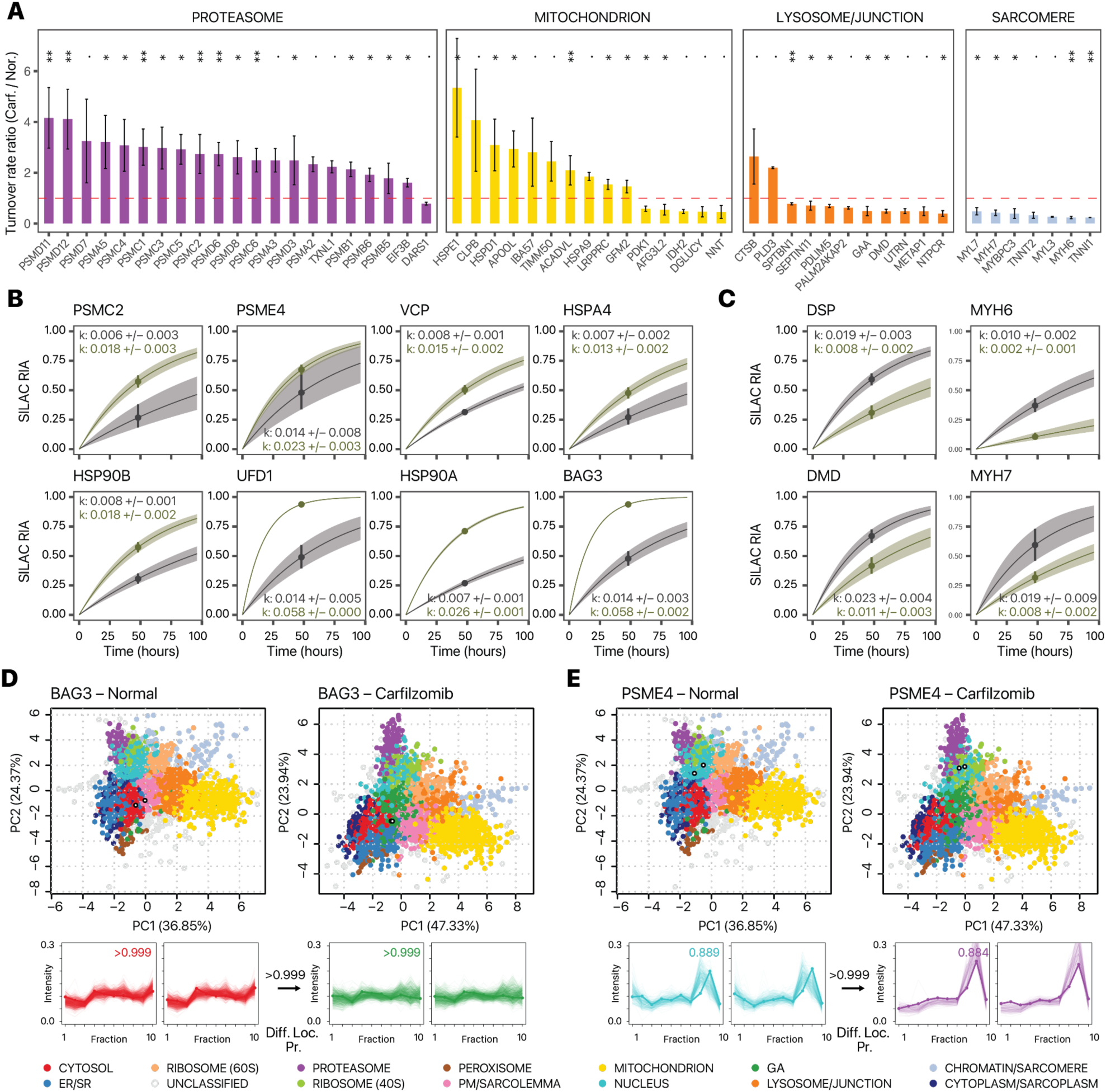
Proteostatic pathways and lesions in carfilzomib mediated cardiotoxicity in iPSC-CMs. **A.** Changes in protein turnover rates between carfilzomib vs. normal iPSC-CMs across selected cellular compartments; **: P < 0. 01; *: P < 0.05; .: P < 0.1; Mann-Whitney test FDR adjusted P values. error bars: standard error. **B.** Kinetic curve representations of proteins with accelerated temporal kinetics in carfilzomib including PSMC2 which corresponds to the ratio in panel A, as well as additional ERAD proteins and chaperones; gray: normal iPSC-CM; green: carfilzomib. **C.** Kinetic curve representations of slowdown of protein kinetics in DSP, DMD, MYH6, and MYH7, corresponding to the ratios in panel A. **D–E.** Spatial map (PC1 vs. PC2) and ultracentrifugation fraction profiles of **D.** BAG3 and **E.** PSME4 in normal and carfilzomib-treated human iPSC-CM, showing a likely differential localisation in conjunction with kinetics changes. White-filled circles: light and heavy BAG3 or PSME4 in each plot. The kinetic curves of BAG3 and PSME4 are in panel B. Numbers at arrows correspond to BANDLE differential localization probability (Diff. Loc. Pr.).

We observed an interconnectivity of spatial and temporal changes, with 23 out of 339 pairs of confident translocators also showing significant kinetic changes. BAG3, a muscle chaperone important for sarcomere turnover (Martin et al., 2021a), shows elevated kinetics (**Figure 7B**) and a partition away from the soluble cytosol compartment (**Figure 7D**) toward an expanded compartment in carfilzomib that co-sediments with Golgi markers. Inspection of existing annotations show that the majority of categorized proteins are not canonical Golgi proteins but contain cytoplasm and endosome terms; hence it likely represents a less soluble cytoplasmic fraction consistent with a lower abundance in the last ultracentrifugation step (**Supplemental Figure S13**, **Supplemental Figure S14, Supplemental Data S9**). This is consistent with the known dynamic partitioning of BAG3 between the cytosol and myofilament fractions for its function (Martin et al., 2021b). Secondly, we find that accelerated temporal kinetics of PA200/PSME4 proteasome activator (**Figure 7B**) in conjunction with a change in localization from the nuclear compartment in baseline toward the proteasome compartment upon carfilzomib (**Figure 7E**). The PSME4/PA200 proteasome activator is known to bind with the 20S/26S proteasome complex to stimulate proteolysis and has a putative nuclear localization signal (Ustrell et al., 2002). The change in localization is therefore consistent with increased binding with the proteasome complex. In parallel, the proteasome activator PA28/PSME3 also relocalizes to the proteasome upon carfilzomib (**Supplemental Figure S15**), altogether suggesting a remodeling of proteasome configuration upon carfilzomib.

Taken together, the spatiotemporal proteomics data here identified major proteostatic pathways induced in carfilzomib, involving a potential remodeling of the proteasome, induction of chaperones and ERAD proteins, and mitochondrial protein quality control mechanisms that may be important for preserving function. On the other hand, a preferential decrease of temporal kinetics in sarcomere and desmosome proteins suggest that the interruption of protein quality control and turnover in these important cardiomyocyte components may be principal sites of lesion in carfilzomib cardiotoxicity. Finally, we assessed the protein-level expression profiles in the hearts of mice treated with carfilzomib for 2 weeks to model cardiac dysfunction (Supplemental Methods). Notably, we find differential protein abundance analysis showed that MHC-β (MYH7) and desmoplakin (DSP) are the 1st and 5th most significantly up-regulated proteins among 3,379 quantified proteins in the hearts of mice treated with carfilzomib (**Supplemental Figure S16**), consistent with their accumulation following proteasome inhibition and suggesting the possibility that similar proteostatic lesions may underlie cardiotoxicity mechanism in vivo.

## Discussion

Advances in spatial proteomics have opened new avenues to discover the subcellular localization of proteins on a proteome scale. Thus far however, few efforts have linked the spatial dynamics of the proteome to other dynamic proteome parameters, which hinders a multi-dimensional view of protein function (Burnum-Johnson et al., 2022; Larance et al., 2013). Co-labeling of SILAC (MS1) and TMT (MS2) tags have been used to increase the channel capacity of quantitative proteomics experiments (Dephoure and Gygi, 2012). Here we adopted the extended labeling capacity to encode spatial and temporal information in the same experiment (**Figure 1D**). The SPLAT design has the advantages of resolving proteome-wide temporal kinetics and spatial distributions in the same experiments and identifying their interactions such as compartment-specific turnover; secondly, it allows separate observations of spatial distribution of new vs. old protein pools. Despite each of the SILAC labeled pairs (light and heavy) and their associated TMT profiles being separately quantified by mass spectrometry as independent ions, the spatial profiles of light and heavy proteins are highly similar under baseline conditions (**Figure 4A-B**) which provides additional assurance of spatial assignments.

Applying the workflow to human AC16 cardiac cells under thapsigargin and tunicamycin induced ER stress and the associated UPR, we observed that both ER stress inducing drugs led to a global suppression in turnover rate, consistent with the reduced translation known to be caused by ER stress. The temporal kinetics data of individual proteins revealed the coordinated activation of known and suspected stress mediators through their increased kinetics, particularly concentrated in the ER and Golgi compartments (**Figure 2E**). At the same time, the spatial proteomics profiles revealed substantial endomembrane remodeling and hundreds of translocating proteins under ER stress. A recent work also reported ∼75 translocator protein candidates under acute low-dose thapsigargin (250 nM for 1 hr) in U-2 OS cells, although the experimental approach was more optimized toward mRNA detection (Villanueva et al., 2022). The spatial data here add to an emerging view of dynamical protein regulation under cellular stress, illustrating a differential localization of RNA binding proteins to stress granules, targeting of specific proteins toward lysosomes, as well as membrane trafficking of ion channels and amino acid transporters. UPR activation is known to induce the biosynthesis of non-essential or partly-essential amino acids despite protein synthesis suppression (Gonen et al., 2019); the recycling of lysosomal lysine and arginine regulates the sensitivity to ER stress (Higuchi-Sanabria et al., 2020); whereas deprivation of amino acids is known to activate downstream pathways of UPR including CHOP in vitro (Harding et al., 2003). Moreover, the knockdown of SLC3A2 has been found to suppress the activation of ER stress response pathways including ATF4/6 induction (Liu et al., 2018), together suggesting amino acid transporters may function in ER stress response. The data here indicate that these transporters may in turn be regulated by their spatial localization beyond steady-state abundance. Other changes are found that may be specific to stressors. Tunicamycin treatment leads to a further decrease in turnover of lysosomal proteins compared to other organelles. The lysosome plays important roles in recycling of glycans, a process which may be slowed under ER stress in response to the decrease in glycosylation.

Changes in protein spatial distribution can occur due to a relocation of an existing protein, where in a protein may respond to a signaling cue such as a post-translational modification status and subsequently migrate to a subcellular location. Alternatively, an alternate localization of newly synthesized proteins can also drive spatial redistribution. By comparing the translocation behaviors of new and old protein pools separately, we were able to observe a partitioning of new and old protein pools including epidermal growth factor receptor (EGFR/ErbB1/HER1). ErbB family proteins are required for both normal heart development and prevention of cardiomyopathies in the adult heart. EGFR is a receptor tyrosine kinase of this family capable of triggering multiple signaling cascades, and can be activated via both ligand dependent and ligand independent pathways. Upon ER stress induction, EGFR immunofluorescence showed a partial translocation away from the plasma membrane. Although immunofluorescence cannot distinguish between old and new protein pools, this partial translocation is consistent with the mass spectrometry data showing partial translocation, involving the newly synthesized heavy protein pool. We hypothesize that this partial translocation is suggestive of a ligand-independent trafficking of newly synthesized protein, rather than ligand dependent activation and internalization that is agnostic to protein lifetime. Of note, ligand-independent activation and internalization of EGFR has been previously induced via both starvation and tyrosine kinase inhibitor treatment, leading to cellular autophagy (Tan et al., 2015), hence this partial translocation may carry functional significance to protective cellular response. We further applied SPLAT to a different, non-proliferating cell type, namely human iPSC-derived cardiomyocytes, which have gained increasing utility for modeling the cardiotoxic effects of the cancer drugs, including carfilzomib. In the carfilzomib experiment, the data from SPLAT revealed a surprising similarity in global turnover rates between control and treatment. These observations are consistent with a compensatory rescue of proteasome abundance and activity previously observed in proteasome inhibition by carfilzomib (Demo et al., 2007; Forghani et al., 2021) or bortezomib (Meiners et al., 2003) in other cell types. Proteasomes are known to be regulated by negative feedback mechanisms (Meiners et al., 2003; Xie and Varshavsky, 2001; Xu et al., 2008), which could explain the lack of change in proteome-wide half-life differences and instead suggest that toxicity may derive from more specific cellular lesions. We identified a significant reduction in turnover in sarcomere proteins, which may be particularly sensitive to interruptions in proteasome activity and moreover may account for the bulk of turnover flux in iPSC-CMs given their high abundance. Finally, the activation and translocation of proteostatic pathway proteins BAG3 and PA200/PSME4 in cardiac cells may be explored as potential targets to ameliorate proteostatic disruptions and cardiotoxic effects.

### Limitations

Although there is no inherent limit to the number of dynamic SILAC labeling time points that can be investigated, in our experiments we have used only a single time point per treatment (16 hours post thapsigargin or tunicamycin; 48 hours post carfilzomib), which was selected based on the drug treatment models but also needed to be sufficient to capture the median half-life of proteins in the cell types studied (**Figure 2D**; **Figure 6J**). Hence the collection time points need to be optimized for different cell types with distinct intrinsic protein turnover rates. Synthesized proteins may have further relocalized following the start of dynamic SILAC labeling, hence some acute translocation responses may be missed. Although the inclusion of earlier time points might provide insight into early translocation events, reaching this objective may be technically challenging, as the acquisition of spatial localization information from the heavy SILAC labeled peptides would be hindered by their low intensity. The double labeling design also requires independent MS2 acquisition of light and heavy peptides, which can decrease the depth and data completeness of mass spectrometry-based proteomics analysis. Future work may alleviate this limitation by modifying the mass spectrometry acquisition methods to automatically trigger the acquisition of heavy peptides and reduce incomplete light-heavy pairs.

Secondly, SPLAT shares limitations that are common to common spatial proteomics strategies. The differential ultracentrifugation method employed here requires ∼10^7^ cells and cannot resolve some subcellular fractions, e.g., lysosome from cell junction. The number of classifiable subcellular localizations here is in line with other LOPIT-DC studies, and may be improved in future work that attempts to couple turnover analysis to gradient-based sedimentation approaches with higher spatial resolution. Protein correlation profiling based techniques generally face challenges in recognizing proteins with multiple localizations or partial translocations. For instance, the multi-functional ERAD protein p97/VCP is known to have multiple subcellular localizations, but its precise subcellular translocation profile is difficult to interpret from the TMT data and is unclassified to any compartment. Because translocation may be sub-stoichiometric, translocated proteins can have lower confidence in classification of location. Hence, other biologically relevant translocators reside in the data that await exploration. In the thapsigargin and tunicamycin experiments, upward of 1,000 candidate translocation patterns were detected with significant differences in localization (99% probability) using BANDLE, but many proteins presented a challenge to clear interpretation of compartments upon manual inspection. Progress in this area may require development of spatial separation methods that combine orthogonal separation principles.

In summary, we describe an experimental workflow and data analysis pipeline that integrates dynamic time-resolved stable isotope labeling kinetic analysis with differential ultracentrifugation-based subcellular proteomics to characterize proteome-wide spatial and temporal changes upon perturbation. This method may be broadly useful for understanding the function and behaviors of proteins inside the cell, and may provide new insight into the mechanisms that regulate protein stability and localization in stress, disease, and drug treatment.

## Methods

Additional methods can be found in the Supplemental Information file.

### AC16 Cell culture, metabolic labeling, UPR induction

AC16 cells procured from Millipore between passage number 11 and 16 were cultured in DMEM/F12 supplemented with 10% FBS and no antibiotics. Cells were maintained at 37°C with 5% CO2 and 10% O2. For isotopic labeling, SILAC DMEM/F12 (Thermo Scientific) deficient in both L-lysine and L-arginine was supplemented with 1% dialyzed FBS and heavy amino acids ^13^C6^15^N2 L-Lysine-2HCl and ^13^C6^15^N4 L-Arginine-HCl (Thermo Scientific) at concentrations of 0.499 mM and 0.699 mM, respectively. Light media was switched to heavy media and cells were labeled for 16 hours prior to harvest. UPR was induced with 1 µM thapsigargin (SelleckChem) or 1 µg/mL tunicamycin (Sigma) at the same time as isotopic labeling.

### Cell harvest, differential centrifugation, and isobaric labeling

Cell harvest and subcellular fractionation was performed based on the LOPIT-DC differential ultracentrifugation protocol as described in Geladaki et al. (Geladaki et al., 2019). Briefly, AC16 cells were treated, harvested with trypsinization, washed 3× with room temperature PBS, and resuspended in a detergent free gentle lysis buffer (0.25 M sucrose, 10 mM HEPES pH 7.5, 2 mM magnesium acetate). 1.5 mL of suspension at a time was lysed using an Isobiotec ball bearing homogenizer with a 16 µM clearance size until ∼80% of cell membranes were lysed, as verified with trypan blue (approximately 15 passes through the chamber). Lysates were spun 3 times each in a 4°C swinging bucket centrifuge 200 × g, 5 min to remove unlysed cells. The supernatant was retained and used to generate the 9 ultracentrifugation pellets using spin parameters shown in **Supplemental Table S2.**

The supernatant generated in the final spin was removed and all pellets and the final supernatant were stored at -80°C until proceeding. Supernatant was thawed on ice and precipitated in 3× the volume of cold acetone overnight at –20°C.This was used to generate pellet 10 by centrifuging at 13,000 × g for 10 minutes at 4°C. Excess acetone was removed and the pellet was allowed to dry briefly before resuspension in a resolubilization buffer of 8 M urea, 50 mM HEPES pH 8.5, and 0.5% SDS with 1x Halt Protease and Phosphatase Inhibitor Cocktail (Thermo Scientific). The suspension was sonicated in a Biorupter with settings 20× 30s on 30s off at 4°C.

Pellets from the ultracentrifugation fractions 1 to 9 were resuspended in RIPA buffer with Halt Protease and Phosphatase Inhibitor Cocktail (Thermo Scientific) with sonication in a Biorupter with settings 10x 30s on 30s off at 4°C. Insoluble debris was removed from all samples (1-10) by centrifugation at 14,000 × g, 5 minutes. Protein concentration of all samples was measured with Rapid Gold BCA (Thermo Scientific). The samples were digested and isobarically tagged using the iFASP protocol (McDowell et al., 2013). 25 ug protein per sample in 250 uL 8M urea was loaded onto Pierce Protein Concentrators PES, 10K MWCO prewashed with 100 mM TEAB. The samples were again washed with 8 M urea to denature proteins and remove SDS. The samples were washed with 300 uL 100 mM TEAB twice. The samples were then reduced and alkylated with TCEP and CAA for 30 minutes at 37°C in the dark. CAA and TCEP were removed with centrifugation and the samples were washed 3x with 100 mM TEAB. Samples were digested atop the filters overnight at 37°C with mass spectrometry grade trypsin (Promega) at a ratio of 1:50 enzyme:protein. A total of 0.2 mg of TMT-10plex isobaric labels (Thermo Scientific) per differential centrifugation fraction were equilibrated to room temperature and reconstituted in 20 µL LC-MS grade anhydrous acetonitrile. In each experiment, labels were randomly assigned to each fraction (**Supplemental Tables S3-S4**) with a random number generator to mitigate possible batch effect. Isobaric tags were added to peptides still atop the centrifugation filters and incubated at room temperature for 1 hour with shaking. The reactions were quenched with 1 µL 5% hydroxylamine at room temperature for 30 minutes with shaking. Labeled peptides were eluted from the filters with centrifugation. To further elute labeled peptides 40 µL 50 mM TEAB was added and filters were again centrifuged. All 10 labeled fractions per experiment were combined and mixed well before dividing each experiment into two aliquots. Aliquots were dried with speed-vac and stored at –80°C.

### Liquid chromatography and mass spectrometry

One aliquot per experiment was reconstituted in 50 µL 20 mM ammonium formate pH 10 in LC-MS grade water (solvent A) for high pH reverse phase liquid chromatography (RPLC) fractionation. The entire sample was injected into a Jupiter 4 µm Proteo 90 Å LC Column of 150 × 1 mm on a Ultimate 3000 HPLC system. The gradient was run with a flow rate of 0.1 mL/min as follows: 0–30 min: 0%–40% Solvent B (20 mM ammonium formate pH 10 in 80% LC-MS grade acetonitrile); 30–40 min: 40%-80% Solvent B; 40–50 min: 80% Solvent B. Fractions were collected every minute and pooled into a total of 20 peptide fractions, then dried with speed-vac.

The dried fractions were reconstituted in 10 µL each of pH 2 MS solvent A (0.1% formic acid) and analyzed with LC-MS/MS on a Q-Exactive HF orbitrap mass spectrometer coupled to an LC with electrospray ionization source. Peptides were separated with a PepMap RSLC C18 column 75 µm x 15 cm, 3 µm particle size (ThermoScientific) with a 90 minute gradient from 0 to 100% pH 2 MS solvent B (0.1% formic acid in 80% LC-MS grade acetonitrile). Full MS scans were acquired with a 60,000 resolution. A stepped collision energy of 27, 30 and 32 was used and MS2 scans were acquired with a 60,000 resolution and an isolation window of 0.7 m/z.

### Mass spectrometry data processing and turnover analysis

Mass spectrometry raw data were converted to mzML format using ThermoRawFileParser v.1.2.0 (Hulstaert et al., 2020) then searched against the UniProt Swiss-Prot human canonical and isoform protein sequence database (retrieved 2022-10-27) using Comet v.2020_01_rev3 (Eng et al., 2015). The fasta database was further appended with contaminant proteins using Philosopher v4.4.0 (total 42,402 forward entries). The search settings were as follows: peptide mass tolerance: 10 ppm; isotope error: 0/1/2/3; number of enzyme termini: 1; allowed missed cleavages: 2; fragment bin tolerance: 0.02; fragment bin offset: 0; variable modifications: TMT-10plex tag +229.1629 for TMT experiments, and lysine + 8.0142, arginine + 10.0083 for all SILAC experiments; fixed modifications: cysteine + 57.0214. The search results were further reranked and filtered using Percolator v3.0 (The et al., 2016) at a 5% FDR. Following database search, the mzML files and Percolator PSMs were input to the SPLAT pipeline. The dynamic SILAC data were analyzed using RIANA v0.7.1 (Hammond et al., 2022) to integrate the peak intensity within a 25 ppm error of the light (+0), heavy (+8, +10), and double K/R (+16, +18, +20) peptide peaks over a 20 second retention time window encompassing the first and last MS2 scan where the peptide is confidently identified. We then calculated the fractional synthesis of all K/R containing peptides as the intensity of the 0th isotopomer peak (m0) over the sum of applicable light and heavy isotopomers (e.g., m0/m0+m8 for a peptide with one lysine). RIANA then performs intensity-weighted least-square curve-fitting using the scipy optimize function to a first-order exponential rise model to find the best-fit peptide turnover rate. Protein turnover rates are calculated as the harmonic mean of peptide turnover rates

### Subcellular localization classification

Subcellular localization classification and translocation predictions were performed using the pRoloc (Gatto et al., 2014) and the BANDLE (Crook et al., 2022) packages in R/Bioconductor. Three replicate batches of AC16 cells per condition each were individually labeled and treated, fractionated and analyzed by mass spectrometry, and biological replicate data were used for pRoloc and BANDLE analysis. Briefly, the subcellular localization markers were selected from the intersecting proteins with a prior data set generated from human U-2 OS osteosarcoma cells (Geladaki et al., 2019) with further curation to account for cell type specific marker expression (**Supplemental Data S10**). A random walk algorithm is used to prune the markers to maximize normalized between-class separation. For differential localization analyses we used the Markov-chain Monte-Carlo (MCMC) and non-parametric model in BANDLE to find unknown protein classification and evaluate differential localization probability. MCMC parameters are 9 chains, 10,000 iterations, and 5,000 burn-in, 20 thinning, seed 42; convergence of the Markov chains is assessed visually by rank plots as recommended (Crook et al., 2019). For additional analysis to describe baseline protein classification and compare MS2 and MS3 performance, a T-augmented Gaussian mixture model with a maximum a posteriori (MAP) method in pRoloc was used.

### Human iPSC-derived cardiomyocytes and proteasome inhibition

Human AICS-0052-003 induced pluripotent stem cell (iPSC) (mono-allelic C-terminus mEGFP-tagged MYL7 WTC; Allen Institute Cell Collection) line was acquired from Coriell Institute and seeded onto Geltrex (Gibco) coated 6 well plates and maintained in StemFlex (Thermo Scientific) media at 37 °C, 5% CO2 with daily media changes. At 80% confluency, cells were passaged using 0.5 mM EDTA before resuspension in StemFlex supplemented with 10 µM Y-27632 (Selleck). Cells were replated into Geltrex coated 12 well plates at a density of 3 × 10^5^ cells/well and daily media changes of StemFlex continued until the cells reached 80% confluency, day 0 of cardiac differentiation. The iPSCs were differentiated into cardiomyocytes using a small molecule based GSK-3 inhibition-Wnt inhibition protocol (Burridge et al., 2014). Briefly, on day 0, cell media was replaced with 2 mL/well RPMI supplemented with B-27 minus insulin (Gibco) and 6 μM CHIR99021 (STEMCELL); on day 2, the media was changed to 2 mL/well RPMI+B-27 minus insulin. On day 3, the media was changed to 2 mL/well RPMI+B27 minus insulin supplemented with 5 μM IWR-1-Endo (STEMCELL). On day 7, the media was changed to 2 mL RPMI+B27 with insulin. Differentiation was confirmed via visualization of morphology, spontaneous contraction of cells, and imaging of the GFP tagged MYL7/MLC-2a. On day 9, the media was changed to RPMI+B27 with insulin without glucose to select for cardiomyocytes. The cardiomyocytes were then passaged at low density with 2 µM CHIR-99021 (Maas et al., 2021). At approximately 75% confluency on passage 2, CHIR supplemented media was removed and replaced with RPMI B-27 with insulin, and used for experiments on day 25–30 post differentiation. For isotopic labeling, RPMI (Thermo Scientific) deficient in both L-lysine and L-arginine was supplemented with B27 with insulin and heavy amino acids ^13^C6^15^N2 L-Lysine-2HCl and ^13^C6^15^N4 L-Arginine-HCl (Thermo Scientific) at concentrations 0.219 mM and 1.149 mM, respectively. 48 hours after CHIR removal, light media was replaced with this heavy media and cells were labeled for 48 hours prior to harvest. 0.5 µM carfilzomib (Selleck) was added with heavy media in the treatment group. Harvesting and ultracentrifugation proceeded as above with the following exception. Due to the diffuse nature of pellets generated in the iPSC-derived cardiomyocytes (iPSC-CM) control experiment, the MLA-50 (Beckman) rotor was switched to the TLA-55 rotor after generation of pellet 5. Consistent force (g) of each spin was maintained by increasing the smaller rotor’s RPM on subsequent spins. This change was repeated during the iPSC-CM treatment experiment. Proteins from each cellular fraction were digested and analyzed with mass spectrometry as above.

## Data Availability

Raw mass spectrometry data have been uploaded to ProteomeXchange under the accession numbers PXD038054, PXD041386, PXD046669, PXD046670, and PXD046671. Source data are provided with this paper.

## Code Availability

Software code for the SPLAT pipeline is available on GitHub at https://github.com/lau-lab/splat, https://github.com/ed-lau/riana and https://github.com/ed-lau/pytmt.

## Author Contributions

J.C. and E.L. conceptualized the study. J.C., V.M., S.K.R. V.H., C.L., V.A., R.W.L., K.Z., J.P., J-W.R. and M.P.L. performed experiments. J.C., V.M., and E.L. processed the data and interpreted the results. J.C. and E.L. wrote software code and performed analysis. J.C., S.K.R., V.M., and E.L. drafted and revised the manuscript. J-W.R., Z.V.W., M.P.L., and E.L. managed funding. All authors read and approved the final version of the manuscript.

## Acknowledgments

The authors thank Dr. Christopher Ebmeier (Director, Proteomics and Mass Spectrometry Core Facility, University of Colorado Boulder) for his assistance with TMT-MS3 experiments. This work was supported in part by NIH award K08-HL148540 to J-W.R., NIH awards R01-HL141278 and R01-GM144456 to M.P.L. and NIH awards R35-GM146815 to E.L; and the University of Colorado SOM Translational Research Scholars Program (TSRP) award to E.L..

