## Supplemental Information for "Simultaneous proteome localization and turnover analysis reveals spatiotemporal features of protein homeostasis disruptions"

**Supplemental Methods** **2**
TMT labeling assignment, isotope correction, and purity assessment 2
MS3-based TMT quantification experiment 2
Unfractionated protein abundance measurement 3
Immunofluorescence staining and microscopy 3
Calculation of euclidean distance between fraction profiles 4
Seahorse extracellular metabolic-flux assay 4
Autophagy assay 4
Proteasome/protease inhibition assays 4
Animal husbandry and carfilzomib treatment 5
Echocardiography 5
Mouse heart protein abundance measurements 5
Database annotation 6
**Supplemental Data** **7**
**Supplemental Tables** **7**
**Supplemental Figures** **9**

### Supplemental Methods

**TMT labeling assignment, isotope correction, and purity assessment**

TMT-10 plex lots were #WF309595 for control AC16 replicate 1 and 2; thapsigargin treated AC16 replicate 1, and tunicamycin treated AC16 replicate 1; and #XB318561 for other samples. Isotope impurities in TMT tags can lead to up to 10% spillover to neighbor channels and decrease quantitative accuracy (**Supplemental Table S1**). We used the non-negative least square algorithm in scipy (Virtanen et al., 2020) to solve for the true channel matrix from the observed channel intensity and impurity matrix for downstream quantification. Spectral purity was calculated with Philosopher freequant (da Veiga Leprevost et al., 2020) within FragPipe using the MSFragger (Kong et al., 2017) built-in TMT10 workflow modified to include heavy K (8.0142) and R (10.00827) within the MSFragger variable modifications. mzML files were searched against the UniProt SwissProt human canonical and isoform protein sequence database (retrieved 2023.08.17) appended with decoys and common contaminants.

**MS3-based TMT quantification experiment**

The SILAC-TMT labeled sample (Control Replicate 2) was cleaned up with a HLB Oasis 1cc (10mg) cartridge. Approximately 20 µg multiplexed peptides were fractionated with high pH reversed-phase C18 UPLC using a 0.5 mm X 200 mm custom packed Dr. Maisch C18-AQ 1.9 µm 120Å column with mobile phases 0.1% (v/v) aqueous ammonia, pH10 in water and acetonitrile (ACN). Peptides were gradient eluted at 20 µL/minute from 2 to 50% ACN in 50 minutes concatenating 12 times for 12 fractions using a Waters M-class UPLC (Waters). Peptide fractions were then dried in a speedvac vacuum centrifuge and stored at -20°C until analysis. High pH peptide fractions were suspended in 3% (v/v) ACN, 0.1% (v/v) trifluoroacetic acid (TFA) and approximately 1 µg tryptic peptides were directly injected onto a reversed-phase C18 1.7 µm, 130 Å, 75 mm by 250 mm M-class column (Waters), using a Waters M-class UPLC (Waters). Peptides were eluted at 300 nL/minute with a gradient from 2% to 25% ACN over 125 minutes then to 50% ACN in 10 minutes and detected using an Orbitrap Fusion Tribrid mass spectrometer (Thermo Scientific). Precursor mass spectra (MS1) were acquired at a resolution of 120,000 from 400 to 1600 m/z with Standard automatic gain control (AGC) target and an Auto maximum injection time set by the Tune Application v3.3.2782.34. Precursor peptide ion isolation width for MS2 fragment scans was 1.2 m/z with a 3 second cycle time. All MS2 spectra were acquired in the linear ion trap with CID activation Collision energy of 35%. Standard automatic gain control (AGC) target and an Auto maximum injection time for MS2 spectra was also set by the Tune Application v3.3.2782.34. MS3 spectra were collected in the Orbitrap with resolution 50,000 for up to 10 SPS Precursors. The MS isolation window was 1.3 m/z and the MS2 isolation window was 3 m/z. HCD was employed with a Collision energy of 65% and the scan range was 100-500 m/z. The automatic gain control was set to 300% with an auto maximum injection time. Dynamic exclusion was set for 60 seconds with a mass tolerance of ±10 ppm.

**Unfractionated protein abundance measurement**

To measure the protein abundance changes following thapsigargin and tunicamycin, AC16 cells were cultured as described in the main text methods section, without SILAC reagents. At 80% confluency, the cells were treated with UPR inducing compounds (1 µg/ml tunicamycin or 1 µM thapsigargin). The treated wells were harvested at 8 and 16 hour timepoints. Control wells were harvested at 16 hours. For each drug, 3 control wells and 3 replicates per time point were digested, tagged with TMT 10-plex reagents (Thermo), fractionated with RPLC, and analyzed with LC-MS/MS as in the SPLAT experiments. Database search and quantification was performed as described in the main text methods section, in the absence of variable SILAC modifications.

**Immunofluorescence staining and microscopy**

For imaging, AC16 cells were seeded in chamber slides (Thermo) and cultured as described above. Following treatment cells were fixed in 4% formaldehyde for 15 minutes, permeabilized with 0.5% TritonX-100 for 15 minutes, and blocked with 3% BSA for 1 hour at room temperature. Cells were incubated with primary antibody for 1 hour at room temperature in 1% BSA. Primary antibodies included anti-Sodium Potassium ATPase (1:500, Abcam, ab76020), anti-EGFR (1:1000, Abcam, ab30) anti-LAMP2 (1:200, Abcam, ab25631), anti-CD98/SLC3A2 (1:200, ProteinTech, 15193-1-AP). The slides were washed with PBS and incubated with secondary antibodies for 1 hour at room temperature in the dark. Secondary antibody reporters included Alexa Fluor® 568 (1:1000, Abcam, ab175471) and Alexa Fluor™ 488 (1:1000, Thermo Scientific, ab150113). The slides were washed and mounted with Fluoroshield DAPI containing mounting media (Abcam), cover slipped, and imaged with an EVOS M5000 microscope (Thermo) and an FV-1000 confocal microscope (Olympus). Images were processed and analyzed using CellProfiler v.4.2.5 (McQuin et al., 2018). The mean intensity of the labeled EGFR channel of a 3 pixel border at the cell's edge was divided by mean intensity of the whole cell to estimate translocation of EGFR.

For iPSC-CM imaging, SCVI273 iPSC-CMs maintained on Matrigel-coated (Corning) glass coverslips were fixed with 4% paraformaldehyde (Thermo Scientific) for 10 minutes, permeabilized with 50 µg/mL digitonin in PBS for 10 minutes, and blocked with 1% BSA and 5% serum from the host species of the secondary antibodies for 30 minutes. After fixation, cells were stained according to standard protocols in a PBS buffer containing 0.1% Triton X-100 and 1% BSA with primary antibody dilutions for rabbit anti-cardiac troponin T (Abcam, ab45932) and mouse anti-α-actinin (Abcam, ab18061). Goat anti-rabbit Alexa Fluor™ 488 and goat anti-mouse Alexa Fluor™ 594 (Thermo Scientific) were used as secondary antibodies. After mounting with Prolong GOLD Antifade with DAPI (Thermo Scientific), imaging was performed using a Revolve microscope (ECHO) and processed by ImageJ.

**Calculation of euclidean distance between fraction profiles**

Following column- and row-wise normalization, the distance between light and heavy protein profiles was calculated as their Euclidean distance between an array containing relative abundance across 3 replicate profiles. This distance was divided by the number of replicates. To compare, an equal sample of light – light pairs were randomly selected and calculated as described.

**Seahorse extracellular metabolic-flux assay**

Mitochondrial and glycolytic ATP consumption rates in human SCVI273 iPSC-CMs was measured using the Seahorse Bioscience XF96 Flux Analyzer with Seahorse CF Real-Time ATP Rate Assay Kit (Agilent Technologies) following the manufacturer's instructions. Briefly, cells (30 – 35,000 cells per well) seeded on XF 96-microplates were treated with carfilzomib 0.5 μM for 0, 1, 12, 24, and 48 hours before measurement. After baseline measurements, oligomycin (1 μM) and rotenone/antimycin A (2 μM) were then sequentially added to each well. Basal ATP production rates and oxygen consumption rates (OCR; pmol/min) were measured according to the manufacturer's protocol.

**Autophagy assay**

Autophagy assay was performed using CYTO-ID® autophagy detection kit (ENZ-51031-0050). All steps were performed according to the manufacturer's instructions. In brief, 25-30,000 iPSC-CMs were seeded per well in 96-well plate and recovered for 4 days. The cells were then treated with carfilzomib ( 0.5μM) vs vehicle for 48 hours. The cells were then washed with 1x assay buffer and incubated in the dark at 37°C for 30 min following the addition of a CYTO-ID green detection reagent and Hoechst 33342 nuclear stain. Green detection reagent allows the measurement of autophagolysosome accumulation and Hoechst 33342 stains bright blue heterochromatin foci. After 30 min, cells were washed with 1x assay buffer to remove access dye and analyzed with fluorescence microplate reader (Cytation) using predefined filter sets [green detection reagent: excitation ~480nm, emission ~530nm; Hoechst nuclear stain: excitation ~340nm, emission ~480nm]. Green signal intensity was normalized with blue signal (representing cell numbers) to compare the extent of intracellular autophagic vehicle accumulation.

**Proteasome/protease inhibition assays**

Proteasome activity assay was performed using a Proteasome activity assay kit (Abcam ab107921). All steps were performed according to the manufacturer's instructions. Briefly, iPSCs-CMs (1 million cells/well) were plated onto the 6-well plate and recovered for 3-4 days. The cells were then treated with carfilzomib (0.5 μM) vs vehicle for 48 hours. CMs were homogenized in 0.5% NP-40 in dH2O and samples were further diluted with assay buffer. Then, samples were incubated with an AMC-tagged peptide substrate for 20 min (T1) and then for 30 min at 37° C to produce fluorophore. Finally, the fluorescence was measured by using an ELISA plate reader (Gen5 cytation) at Ex/Em = 350/440. Data were analyzed using graphpad prism and proteasome activity was calculated such that 1 unit of proteasome activity is equivalent to the amount of proteasome activity that generates 1.0 nmol of AMC per minute at 37°C. This experiment was repeated across 3 independent biological and technical replicates.

**Animal husbandry and carfilzomib treatment**

Wild-type C57BL/6N mice were purchased from Charles River Laboratories. All mouse procedures were conducted in accordance with the protocol approved by the Institutional Animal Care and Use Committee (IACUC) of the City of Hope National Medical Center. All mice were maintained in a 12:12 hour light: dark cycle in temperature-controlled rooms with free access to water and chow food. C57BL/6N mice were treated with carfilzomib (dissolved in 5% DMSO + 95% saline, PR-171, Selleckchem) via intraperitoneal (i.p.) injection twice a week at a dose of 8 mg/kg/body weight (Efentakis et al., 2021). Body weight was recorded before every injection. Mice injected with the vehicle (5% DMSO + 95% saline) were used as controls. All treatments were performed for 2 weeks in total.

**Echocardiography**

Cardiac function was evaluated with unconstrained, conscious mice using echocardiography (Visual Sonics, #Vevo 3100, MS400C probe) as previously described (Li et al., 2021). The parasternal long-axis view of B-mode images at the level of left ventricular outflow tract and the parasternal short-axis view of M-mode images at the level of papillary muscles were captured and analyzed to determine various parameters. Heart rate was also recorded. Left ventricular internal diameters at end diastole (LVID, diastolic) and at end systole (LVID, systolic) were measured from M-mode recordings. Fractional shortening = ([LVID, diastolic - LVID, systolic]/LVID, diastolic) × 100%.

**Mouse heart protein abundance measurements**

Left ventricular tissues were resuspended in 1 mL RIPA buffer (Thermo Scientific) supplemented with protease and phosphatase inhibitor (Thermo Scientific). Tissue was homogenized in prefilled tubes containing 2.8 mm ceramic beads using an Omni Bead Ruptor for 20 seconds at speed 5. The homogenized lysate was then subjected to sonication in a Biorupter (Diagenode) with settings 10× 30 sec on 30 sec off at 4°C. Insoluble debris was removed from all samples by centrifugation at 14,000 × g, 5 minutes. Protein concentration of all samples was measured with Rapid Gold BCA. Twenty-five µg of each sample was digested, isobarically tagged, fractionated with RPLC, and analyzed with LC-MS/MS as in the SPLAT experiments. Database search and quantification was performed as above in the absence of variable SILAC modifications.

**Database annotation**

Known stress granule proteins were retrieved from the RNA granule database (Millar et al., 2023). Known subcellular localization were retrieved from UniProt Gene Ontology Cellular Component (CC) terms (UniProt Consortium, 2021) using UniProt.ws (Carlson, 2017) with the following terms: CYTOSOL – cytosol [GO:0005829]; ER – endoplasmic reticulum [GO:0005783] OR endoplasmic reticulum membrane [GO:0005789] OR endoplasmic reticulum lumen [GO:0005788]; GA – Golgi apparatus [GO:0005794] OR Golgi lumen [GO:0005796] OR Golgi membrane [GO:0000139]; LYSOSOME – lysosome [GO:0005764] OR lysosomal membrane [GO:0005765] OR lysosomal lumen [GO:0043202]; MITOCHONDRION – mitochondrion [GO:0005739] OR mitochondrial inner membrane [GO:0005743] OR mitochondrial outer membrane [GO:0005741] OR mitochondrial matrix [GO:0005759] OR mitochondrial respirasome [GO:0005746]; NUCLEUS – nucleus [GO:0005634] OR chromatin [GO:0000785] OR nucleoplasm [GO:0005654] OR nucleolus [GO:0005730]; PEROXISOME – peroxisome [GO:0005777] OR peroxisomal matrix [GO:0005782] OR peroxisomal membrane [GO:0005778]; PM – plasma membrane [GO:0005886] OR cell surface [GO:0009986]; PROTEASOME – proteasome complex [GO:0000502] OR proteasome accessory complex [GO:0022624] OR proteasome regulatory particle [GO:0005838]; RIBOSOME – ribosome [GO:0005840] OR cytosolic ribosome [GO:0022626]; CHROMATIN – chromosome [GO:0005694] OR chromatin [GO:0000785] OR nucleosome [GO:0000786] OR euchromatin [GO:0000791] OR heterochromatin [GO:0000792]"; CYTOPLASM – cytoplasm [GO:0005737] OR cytoskeleton [GO:0005856] OR actin cytoskeleton [GO:0015629] OR microtubule [GO:0005874] OR microtubule cytoskeleton [GO:0015630] OR cortical actin cytoskeleton [GO:0030864] OR actin filament [GO:0005884] OR cortical cytoskeleton [GO:0030863] OR intermediate filament cytoskeleton [GO:0045111].

For iPSC-CM, the RIBOSOME (40S) compartment was matched against ribosome [GO:0005840] OR cytosolic ribosome [GO:0022626] OR eukaryotic 43S preinitiation complex [GO:0016282] OR eukaryotic 48S preinitiation complex [GO:0033290]; the RIBOSOME (60S) compartment was matched against ribosome [GO:0005840] OR cytosolic ribosome [GO:0022626] OR polysomal ribosome [GO:0042788]. The LYSOSOME/JUNCTION compartment was additionally matched against cell-cell junction [GO:0005911] OR adherens junction [GO:0005912] OR catenin complex [GO:0016342] in addition to the LYSOSOME terms above. The CHROMATIN/SARCOMERE compartment was additionally matched against sarcomere [GO:0030017] OR Z disc [GO:0030018] OR muscle myosin complex [GO:0005859] in addition to the CHROMATIN terms above.

### Supplemental Data

Supplemental Data S1 - Abundance changes in ER stress vs. Normal AC16 cells
Supplemental Data S2 - Protein localization and assignment in Normal AC16 cell
Supplemental Data S3 - Turnover rate ratios in Thapsigargin vs. Normal AC16 cells
Supplemental Data S4 - Protein localization and assignment in Thapsigargin AC16
Supplemental Data S5 - Turnover rate ratios in Tunicamycin vs. Normal AC16 cells
Supplemental Data S6 - Protein localization and assignment in Tunicamycin AC16
Supplemental Data S7 - Protein localization and assignment in Normal iPSC-CM
Supplemental Data S8 - Turnover rate ratios in carfilzomib vs. Normal iPSC-CMs
Supplemental Data S9 - Protein localization and assignment in Carfilzomib iPSC-CMs
Supplemental Data S10 - List of canonical compartment markers in spatial experiments

### Supplemental Tables

**Supplemental Table S1: Isotopic Contaminant Matrix of $TMT^{10}$ Lots XB318561 and WF309595**

| Mass Tag | -2 (XB318561) | -2 (WF309595) | -1 (XB318561) | -1 (WF309595) | +1 (XB318561) | +1 (WF309595) | +2 (XB318561) | +2 (WF309595) |
|---|---|---|---|---|---|---|---|---|
| t126 | 0.0% | 0.0% | 0.0% | 0.0% | 7.4% (127C) | 7.4% (127C) | 0.0% (128C) | 0.0% (128C) |
| t127N | 0.0% | 0.0% | 0.1% | 0.0% | 7.8% (128N) | 7.2% (128N) | 0.1% (129N) | 0.0% (129N) |
| t127C | 0.0% | 0.0% | 0.8% (126) | 0.8% (126) | 6.9% (128C) | 6.6% (128C) | 0.1% (129C) | 0.0% (129C) |
| t128N | 0.0% | 0.0% | 1.2% (127N) | 1.2% (127N) | 6.3% (129N) | 6.3% (129N) | 0.0% (130N) | 0.0% (130N) |
| t128C | 0.0% (126) | 0.0% (126) | 1.5% (127C) | 1.3% (127C) | 6.2% (129C) | 5.7% (129C) | 0.2% (130C) | 0.1% (130C) |
| t129N | 0.0% (127N) | 0.0% (127N) | 1.5% (128N) | 1.6% (128N) | 5.7% (130N) | 5.4% (130N) | 0.1% (131) | 1.3% (131) |
| t129C | 0.0% (127C) | 0.3% (127C) | 2.6% (128C) | 2.7% (128C) | 4.8% (130C) | 4.8% (130C) | 0.0% | 0.0% |
| t130N | 0.0% (128N) | 0.0% (128N) | 2.2% (129N) | 2.2% (129N) | 4.6% (131) | 4.6% (131) | 0.0% | 0.0% |
| t130C | 0.0% (128C) | 0.0% (128C) | 3.1% (129C) | 3.1% (129C) | 3.6% | 3.6% | 0.0% | 0.0% |
| t131 | 0.0% (129N) | 0.0% (129N) | 8.7% (130N) | 8.7% (130N) | 3.4% | 3.4% | 0.0% | 0.0% |

**Supplemental Table S2: Centrifugation Speeds and Times Associated with Each Fraction**

| Fraction | Centrifugation Speed (g) | Centrifugation Time (Minutes) |
|---|---|---|
| 1 | 1000 | 10 |
| 2 | 3000 | 10 |
| 3 | 5000 | 10 |
| 4 | 9000 | 15 |
| 5 | 12,000 | 15 |
| 6 | 15,000 | 15 |

| | | |
|---|---|---|
| 7 | 30,000 | 20 |
| 8 | 79,000 | 43 |
| 9 | 120,000 | 45 |
| 10 | 13,000 (Following Precipitation Procedure) | 10 |

**Supplemental Table S3: Fraction Association of Each TMT Tag in Control and UPR AC16 Cells**

| TMT Label | Normal Replicate 1 | Normal Replicate 2 | Normal Replicate 3 | Thapsigargin Replicate 1 | Thapsigargin Replicate 2 | Thapsigargin Replicate 3 | Tunicamycin Replicate 1 | Tunicamycin Replicate 2 | Tunicamycin Replicate 3 |
|---|---|---|---|---|---|---|---|---|---|
| 126 | Fraction 10 | Fraction 10 | Fraction 3 | Fraction 1 | Fraction 10 | Fraction 6 | Fraction 2 | Fraction 2 | Fraction 8 |
| 127N | Fraction 2 | Fraction 9 | Fraction 9 | Fraction 4 | Fraction 8 | Fraction 8 | Fraction 10 | Fraction 5 | Fraction 4 |
| 127C | Fraction 5 | Fraction 2 | Fraction 1 | Fraction 5 | Fraction 3 | Fraction 2 | Fraction 4 | Fraction 6 | Fraction 3 |
| 128N | Fraction 1 | Fraction 4 | Fraction 4 | Fraction 2 | Fraction 5 | Fraction 9 | Fraction 1 | Fraction 10 | Fraction 6 |
| 128C | Fraction 8 | Fraction 1 | Fraction 5 | Fraction 8 | Fraction 4 | Fraction 7 | Fraction 9 | Fraction 3 | Fraction 1 |
| 129N | Fraction 7 | Fraction 8 | Fraction 8 | Fraction 7 | Fraction 9 | Fraction 3 | Fraction 5 | Fraction 8 | Fraction 10 |
| 129C | Fraction 6 | Fraction 5 | Fraction 10 | Fraction 9 | Fraction 7 | Fraction 5 | Fraction 6 | Fraction 4 | Fraction 9 |
| 130N | Fraction 4 | Fraction 6 | Fraction 2 | Fraction 10 | Fraction 6 | Fraction 1 | Fraction 3 | Fraction 9 | Fraction 2 |
| 130C | Fraction 9 | Fraction 3 | Fraction 6 | Fraction 6 | Fraction 1 | Fraction 10 | Fraction 7 | Fraction 1 | Fraction 7 |
| 131 | Fraction 3 | Fraction 7 | Fraction 7 | Fraction 3 | Fraction 2 | Fraction 4 | Fraction 8 | Fraction 7 | Fraction 5 |

**Supplemental Table S4: Fraction Association of Each TMT Tag in Control and Carfilzomib Treated iPSC Trials**

| TMT Label | Control iPSC-CM | Carfilzomib Treated iPSC-CM |
|---|---|---|
| 126 | Fraction 3 | Fraction 9 |
| 127N | Fraction 10 | Fraction 1 |
| 127C | Fraction 2 | Fraction 4 |
| 128N | Fraction 5 | Fraction 6 |
| 128C | Fraction 6 | Fraction 2 |
| 129N | Fraction 4 | Fraction 8 |
| 129C | Fraction 9 | Fraction 5 |
| 130N | Fraction 8 | Fraction 10 |
| 130C | Fraction 1 | Fraction 3 |
| 131 | Fraction 7 | Fraction 7 |

### Supplemental Figures

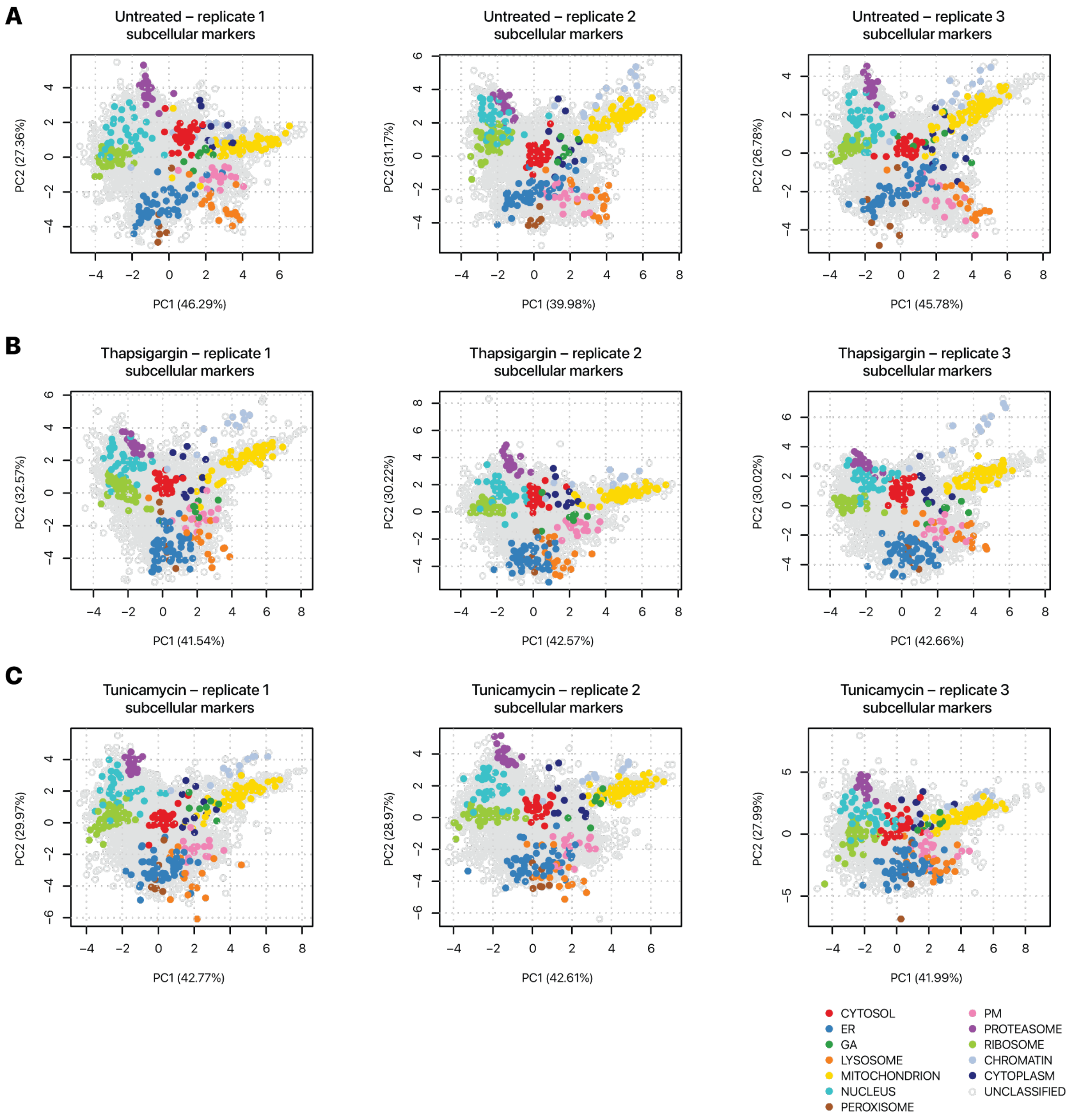


**Supplemental Figure S1**. Separation of subcellular component markers in the spatial proteomics data. Scatter plots of the first (x-axis) and second (y-axis) principal components of ultracentrifugation fraction profiles are shown for each experimental condition: **A.** normal, **B.** thapsigargin, and **C.** tunicamycin treated AC16 cells, n=3 each. Each data point is a protein species. The colored data points correspond to marker proteins known to reside in each of 12 subcellular locations used to train the classification models, showing clear and consistent separation across the experimental conditions. Colors correspond to other spatial maps for AC16 cells throughout the manuscript.

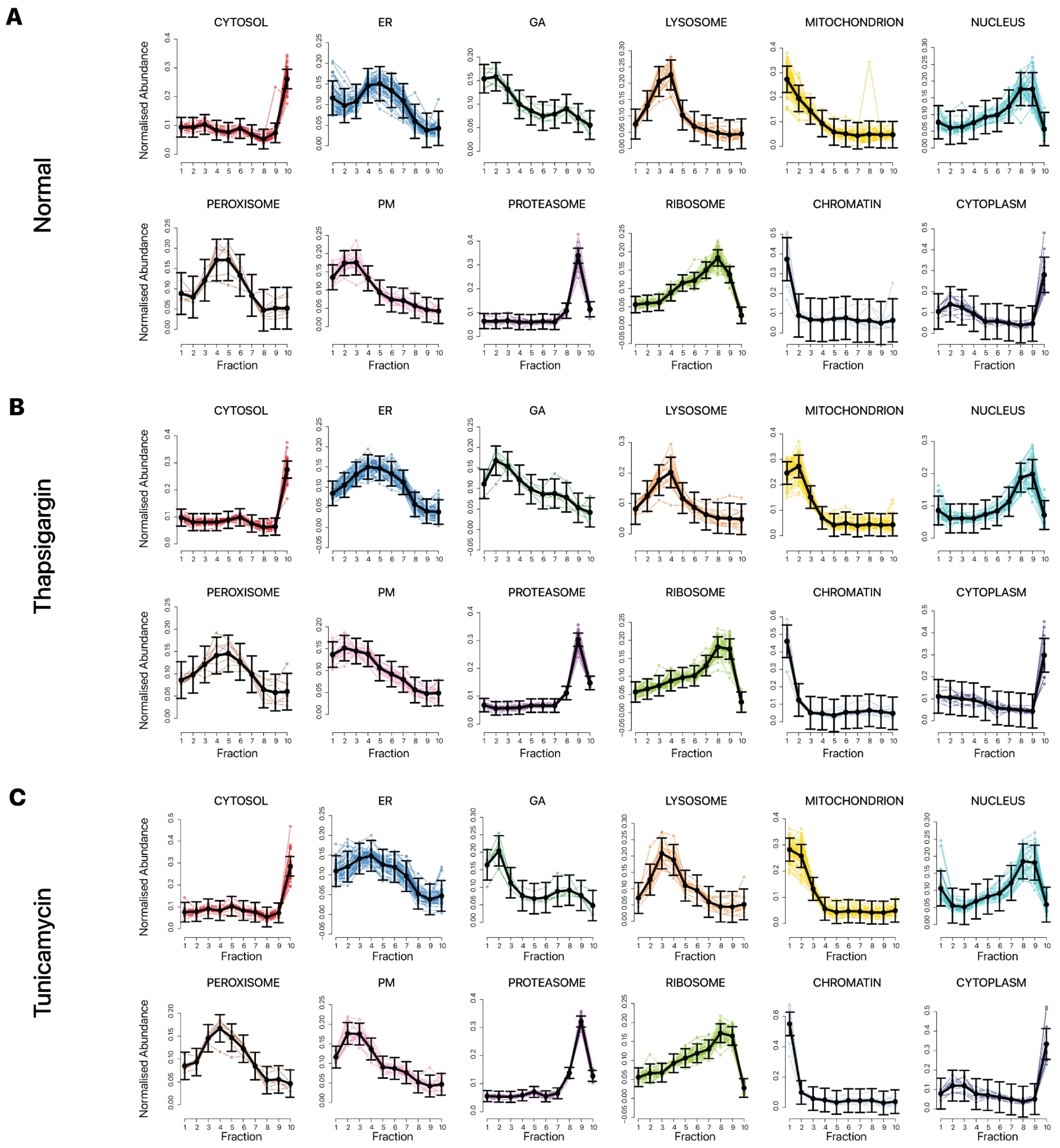


**Supplemental Figure S2.** Ultracentrifugation fraction distributions of cellular component markers. Replicate one of each experimental condition is shown. The line plots show the normalized abundance (y-axis) of marker proteins for each subcellular localization experiment across ultracentrifugation fractions (x-axis) as measured by the TMT channel intensities. The fractions correspond to the ultracentrifugation steps in Supplemental Table S3. Colors correspond to spatial maps for AC16 cells throughout the manuscript. Black lines show average trend line and standard deviation, showing consistent sedimentation profiles of subcellular localization in the **A.** normal, **B.** thapsigargin, and **C.** tunicamycin treated AC16 cells (n=3).

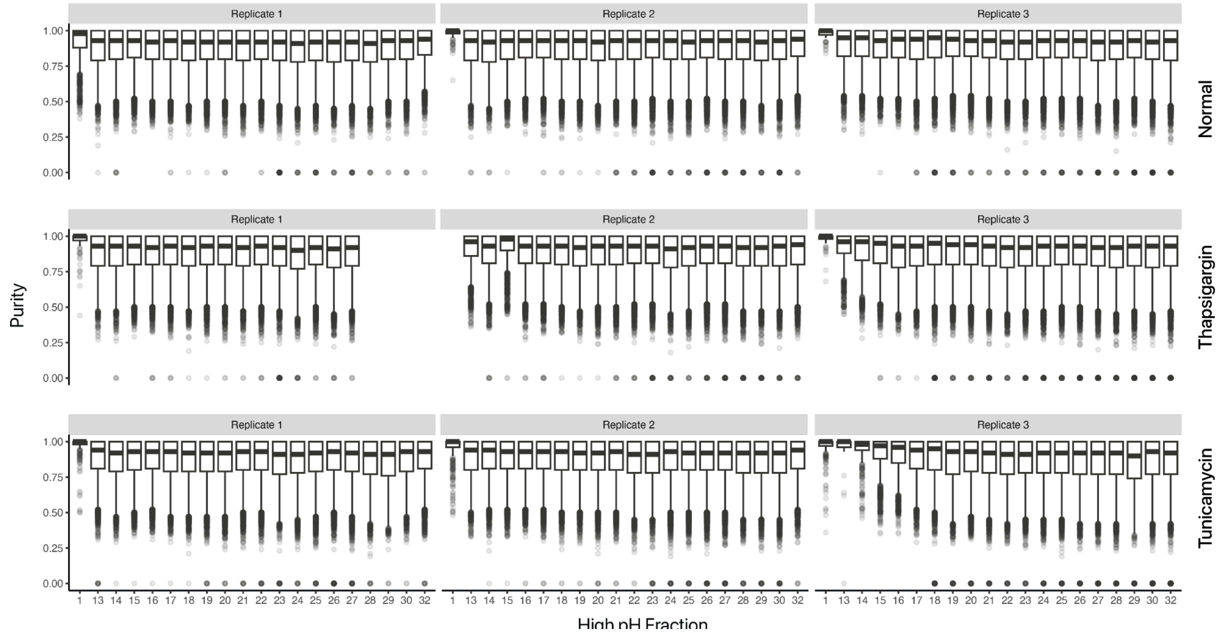


**Supplemental Figure S3.** Spectral purity of MS2-based TMT measurement. Box plots showing the distribution of precursor ion fraction/spectral purity as measured by MSFragger/Philosopher (y-axis) of all confidently identified MS2 scans in the MS2-based TMT measurements across high-pH reversed-phase LC fraction injections (x-axis) in the mass spectrometry experiments in normal, thapsigargin, and tunicamycin treated AC16 cells (n=3), showing high precursor isolation (average 93%) in the MS2 experiment. Center line: median; box limits: interquartile range; whiskers: 1.5x interquartile range; points: outliers.

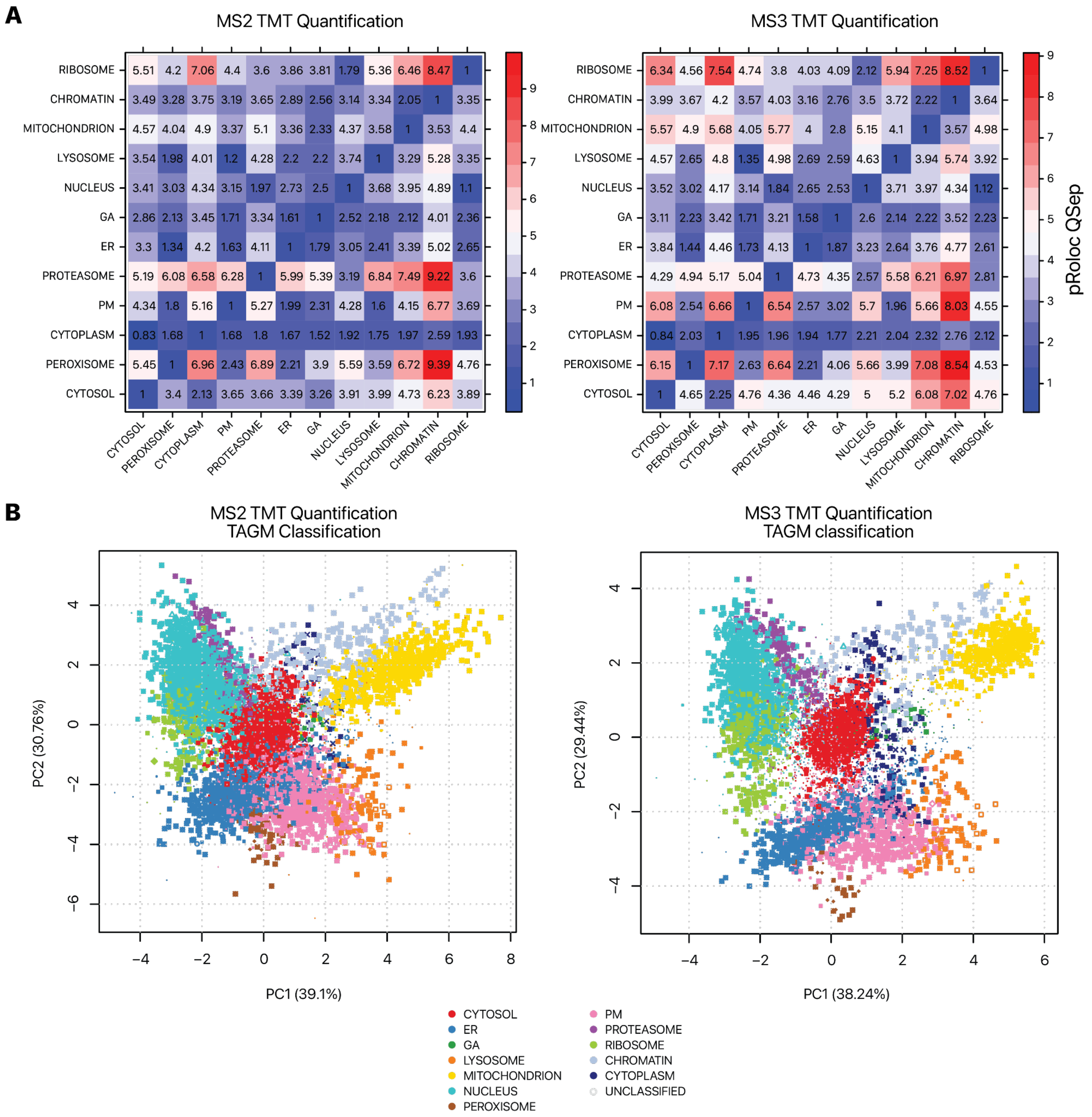


**Supplemental Figure S4.** Comparison of subcellular localization assignment in MS2 and MS3-based TMT measurements. An identical sample (replicate 2 of normal AC16 cells) was analyzed by MS2 and MS3 based quantification. **A.** The QSep index in the pRoloc package reflects the between-cluster distance vs. within-cluster distances of the 12 subcellular locations. MS3 achieved a modest increase in median QSep (3.98 vs. 3.51) suggesting the subcellular component clusters were slightly better separated. **B.** Spatial maps of proteins in MS2 vs. MS3 quantification. Colors correspond to other spatial maps in AC16 cells throughout the manuscript. Data point size reflects the confidence of TAGM-MAP classification.

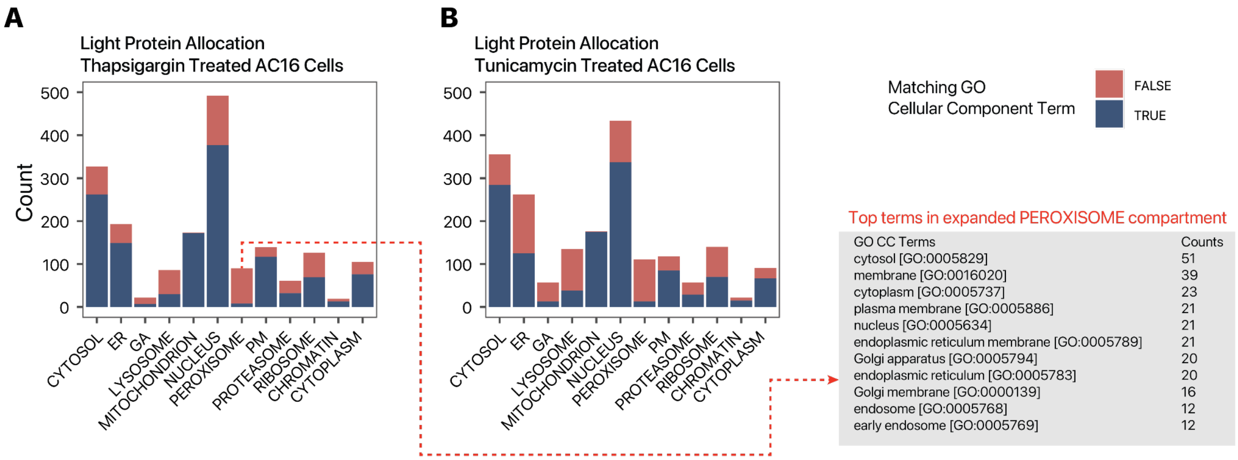


**Supplemental Figure S5.** Correspondence of spatial classification with prior annotations in stressed cells. As related to main Figure 2C, the bar charts show the number of light (i.e., non-heavy-SILAC labeled) proteins (y-axis) classified to each of 12 subcellular locations (x-axis) in thapsigargin (left) and tunicamycin (right) treated AC16 cells (n=3). The colors represent whether proteins classified to each subcellular location are also known to reside in the subcellular component of question in Gene Ontology Cellular Component terms retrieved from UniProt. In normal, thapsigargin, and tunicamycin treated AC16 cells, 69.5%, 71.9%, and 63.0% of classified proteins are consistent with known annotations, respectively; hence the classified subcellular localization match the expected assignments from prior knowledge and are not substantially affected by cellular stressors. The expanded peroxisome compartment in stressed AC16 cells primarily contained non-peroxisome annotated proteins that co-sedimented with the trained peroxisome compartment, and are referred to as the peroxisome/endosome compartment in the manuscript.

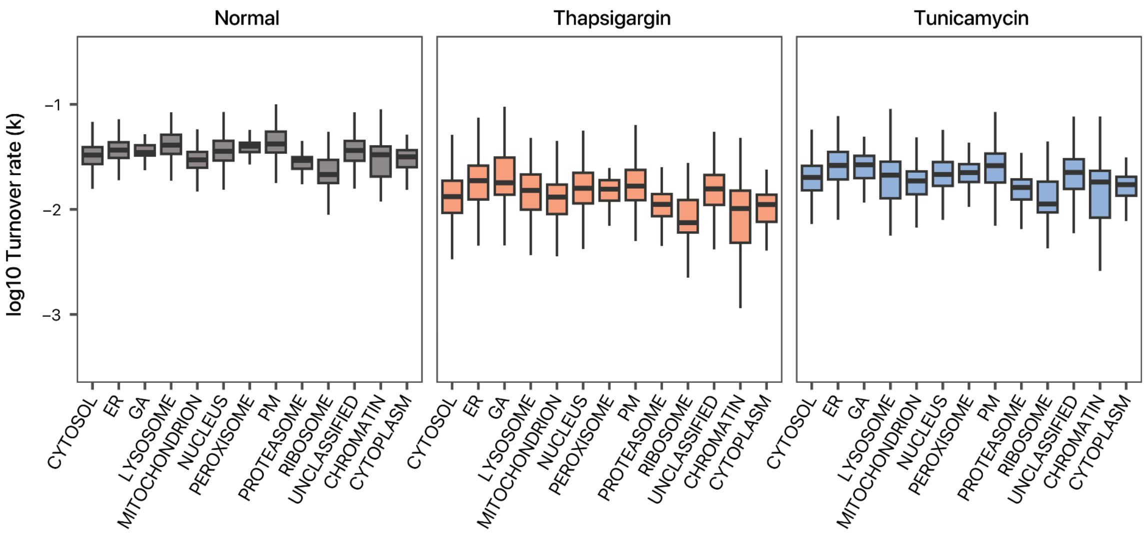


**Supplemental Figure S6.** Subcellular localization differences in protein turnover rate. Boxplots showing the log10 protein turnover rates (k) of proteins assigned to each of 12 subcellular localizations in normal (left), thapsigargin (middle), and tunicamycin (right) treated AC16 cells (n=3). Center line: median; box limits: interquartile range; whiskers: 1.5x interquartile range.

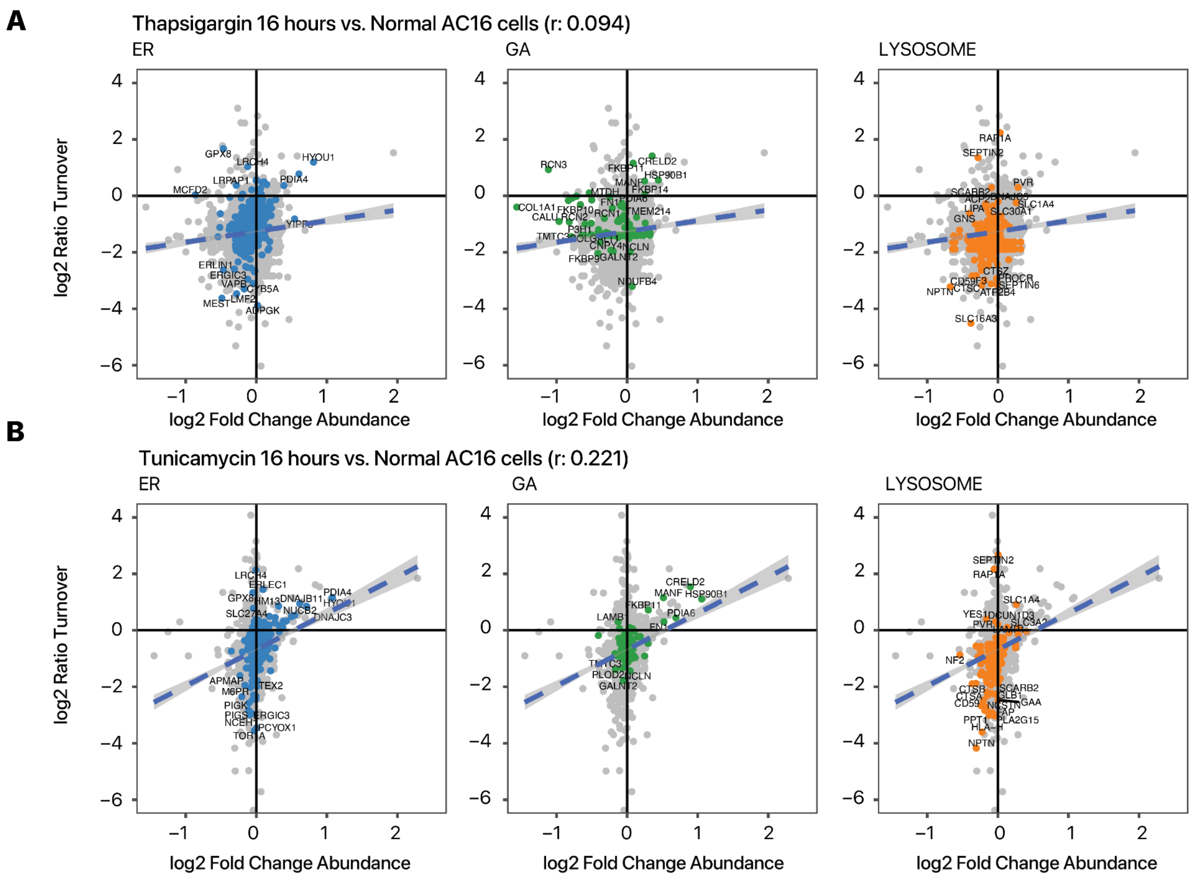


**Supplemental Figure S7.** Protein abundance and turnover changes following thapsigargin and tunicamycin treatment. Scatterplots showing the relationship between log2 of protein abundance fold changes (x-axis) and log2 of turnover ratios (y-axis) in **A.** thapsigargin and **B.** tunicamycin treatment. In each of the series of scatterplots from left to right, proteins assigned to the ER (blue), GA (green), and lysosome (orange) are labeled. Overall protein kinetic changes are only modestly correlated with protein abundance changes.

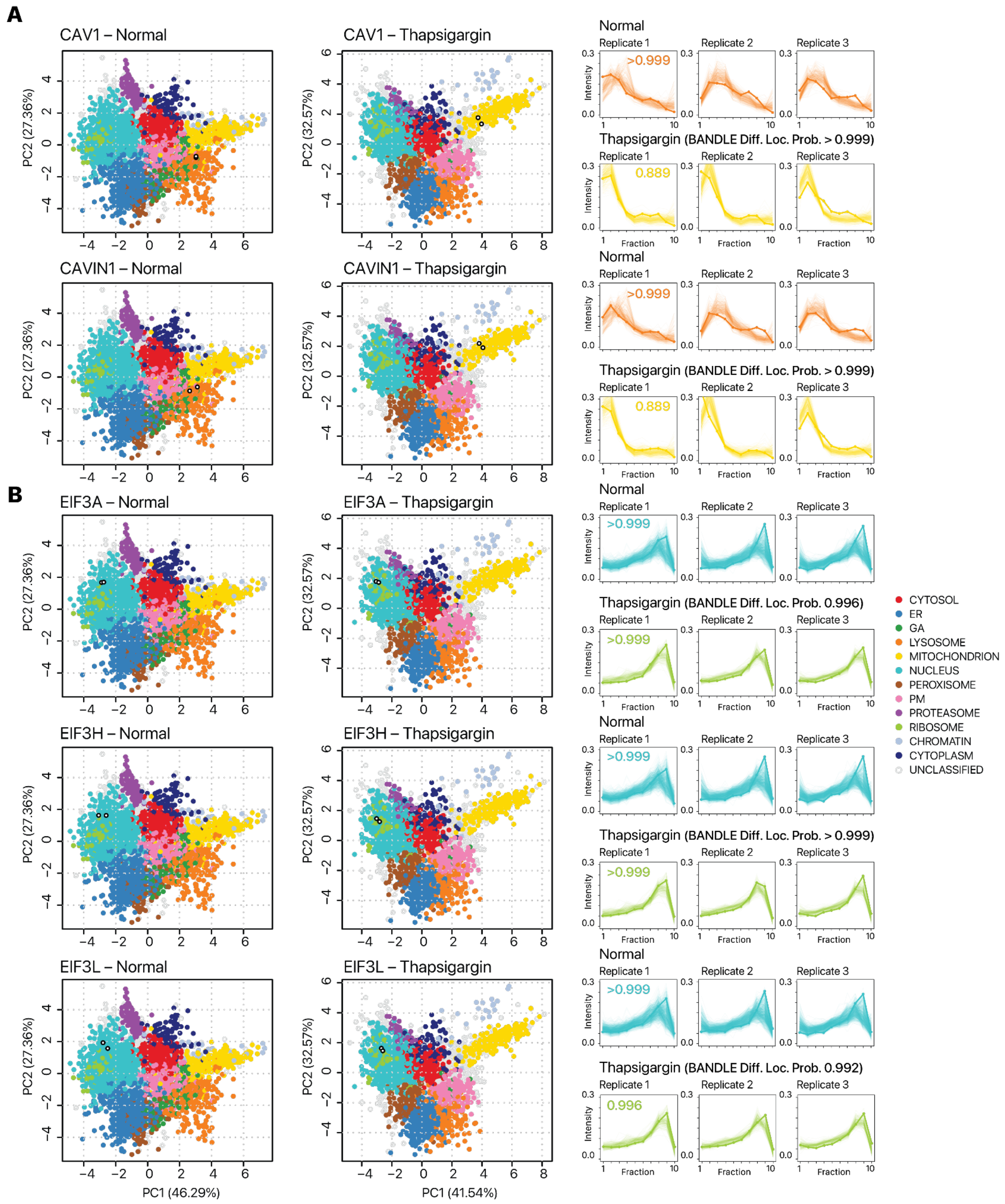


**Supplemental Figure S8.** Known translocation events under cellular stress captured in the spatial proteomics data. **A.** Caveolae migration toward the mitochondrion during cellular stress is reflected in the differential localization of CAV1 and CAVIN1 (BANDLE probability > 0.999) toward the mitochondrion compartment. **B.** A switch to EIF3-dependent translation initiation, a hallmark of prolonged ER stress and integrated stress response, is evident in the differential localization toward the ribosome compartment of three independent EIF3 subunits EIF3A (BANDLE probability: 0.996), EIF3H (BANDLE probability: >0.999), and EIF3L (BANDLE probability: 0.992). The nucleus localization probability of these proteins in normal AC16 cells is accompanied by a high outlier probability (**Supplemental Data S2**) and may reflect partial ribosome localization. Left: spatial maps of PC1 vs PC2; colors correspond to other spatial maps for AC16 cells throughout the manuscript. Open circles: location of light and heavy protein in each condition. Only the map of one of three replicates is shown for simplicity. Right: ultracentrifugation profiles showing relative abundance (y-axis) across fraction (x-axis). Numbers inside the fraction profile correspond to BANDLE localization probability.

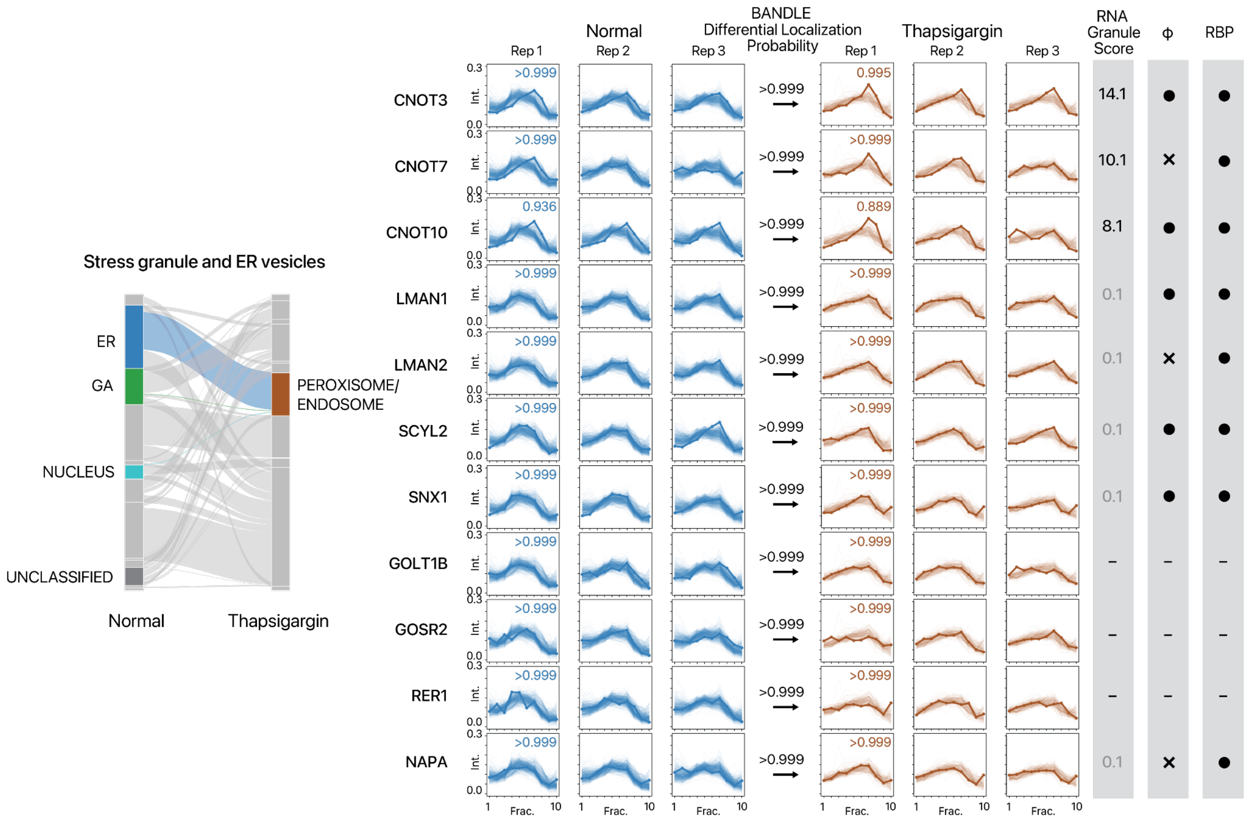


**Supplemental Figure S9.** Additional examples of proteins translocating toward the peroxisome-endosome cosidementing compartment upon thapsigargin treatment. (Left) Alluvial plot of significant protein translocation (Pr > 0.95) from the ER, Golgi apparatus (GA), and nucleus toward the peroxisome. Colors correspond to spatial maps for AC16 cells throughout the manuscript. (Right) Ultracentrifugation fraction profile of CNOT3, CNOT7, CNOT10, LMAN1, LMAN2, SCYL2, SNX1, GOLT1B, GOSR2, RER1, and NAPA showing the localization of the proteins to the ER and to the peroxisome/endosome fraction in normal and thapsigargin-treated AC16 cells, respectively. X-axis: fraction 1 to 10 of ultracentrifugation. Y-axis: relative channel abundance. Bold lines represent the protein of interest; light lines represent ultracentrifugation profiles of all proteins classified to a respective localization. Colors correspond to subcellular localization for all AC16 data throughout the manuscript. Numbers in the box represent BANDLE localization probability to the compartment. RNA Granule Score: score from RNA Granule Database (https://rnagranuledb.lunenfeld.ca/). A score of 7 or above is considered a known stress granule protein. Phi: predicted phase separation participation. RBP: Annotated RNA binding protein on the RNA Granule Database. One circle denotes known RNA binding proteins (RBP) in at least one data set; two circles denote known RBP in multiple datasets.

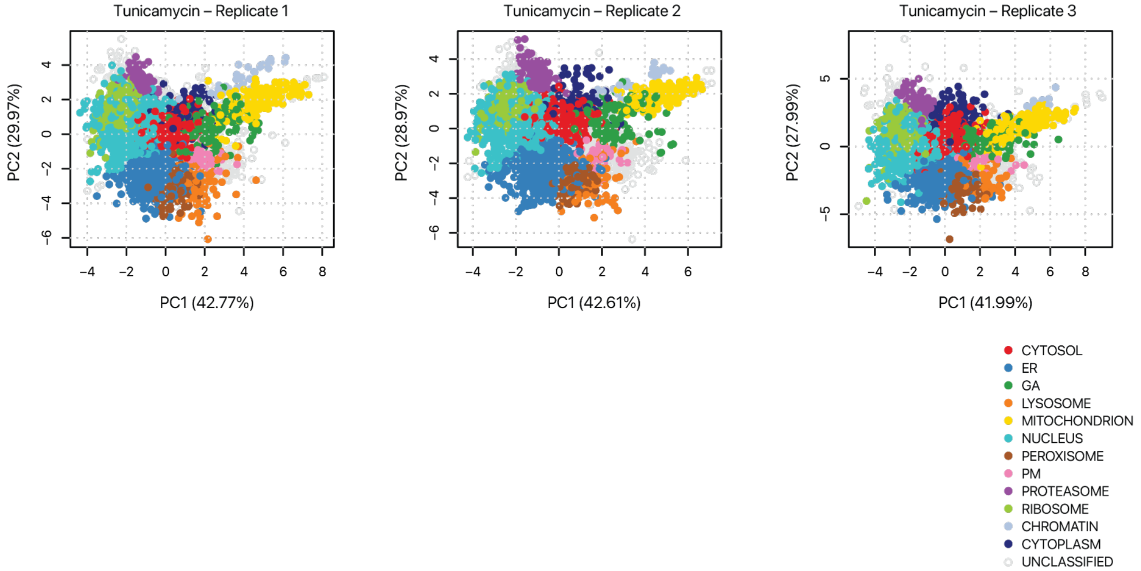


**Supplemental Figure S10:** Subcellular localization of proteins following tunicamycin treatment. PC1 and PC2 of proteins spatial map showing the localization of confidently allocated proteins in tunicamycin-treated AC16 cells. Each data point represents a protein; color represents classification of subcellular localization consistent with other AC16 cell data throughout the manuscript.

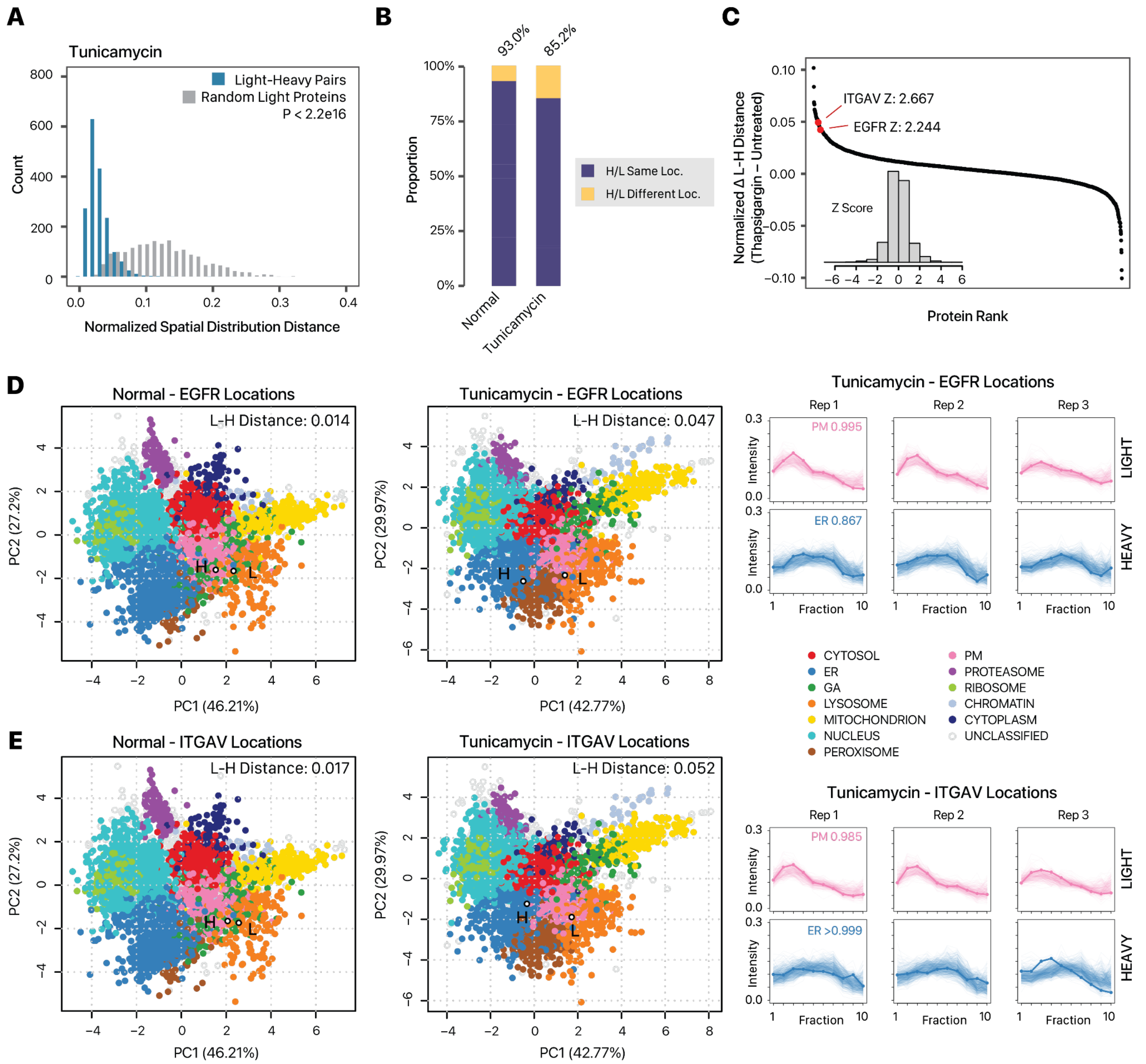


**Supplemental Figure S11:** Partition of new and old EGFR and ITGAV in tunicamycin treatment. **A.** Histogram showing the similarity in light and heavy proteins in normalized spatial distribution distances in tunicamycin-treated AC16 cells. X-axis: euclidean distance of fraction profiles across 3 replicates; y-axis: count. Blue: distance for quantified light-heavy protein pairs. Grey: distribution of each corresponding light protein with another random light protein. P value: Mann-Whitney test. **B.** Proportion of heavy-light protein pairs with confidently assigned localization that are assigned to the same location (purple) in normal (left; 93.0%) and tunicamycin-treated (right; 85.2%) cells. **C.** Ranked changes in heavy-light pair euclidean distance upon tunicamycin treatment. The majority of proteins show no change (+/- 0.02 in euclidean distance). The positions of EGFR and ITGAV are highlighted. Inset: Z score distribution of all changes. The spatial maps for **D.** EGFR and **E.** ITGAV showing a translocation of newly synthesized (heavy; H) but not old (light; L) proteins from the plasma membrane (PM) to the ER fraction in tunicamycin treated AC16 cells. Open circles show the location of the proteins in the map. Numbers denote

BANDLE allocation probability. (Right) Ultracentrifugation profiles showing different sedimentation behaviors of the light and heavy proteins.

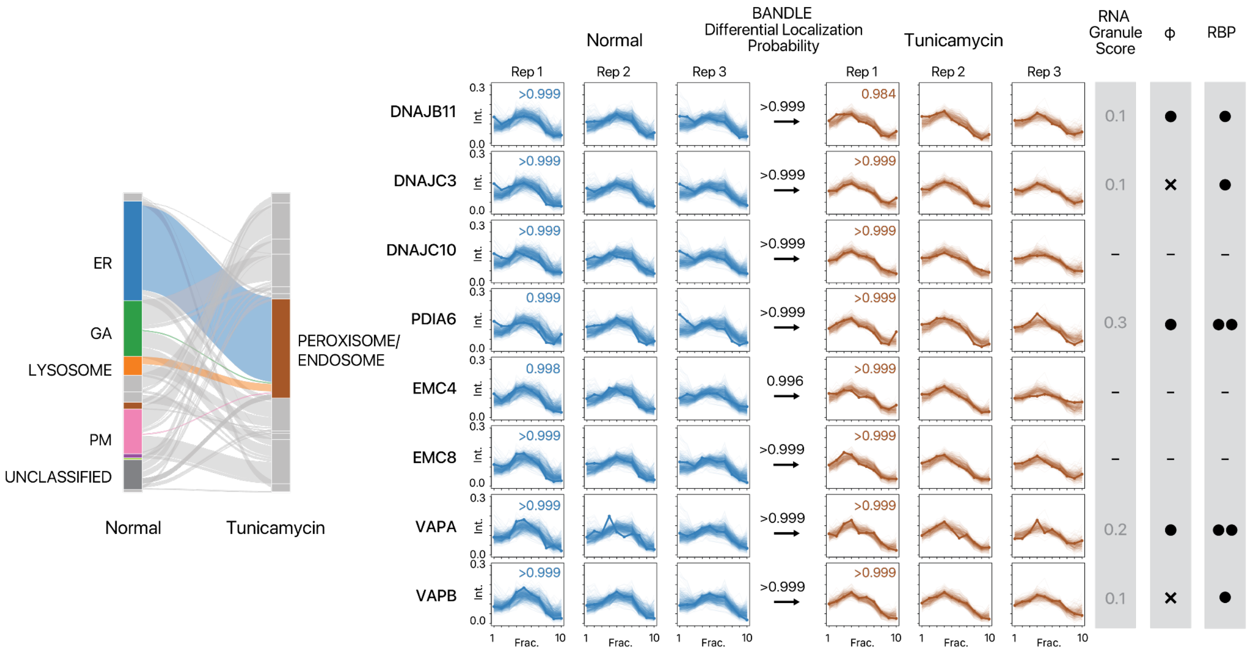


**Supplemental Figure S12.** Examples of proteins translocating toward the peroxisome-endosome cosidementing compartment in tunicamycin treatment. (Left) Alluvial plot of significant protein translocation (Pr > 0.95) from the ER, Golgi apparatus (GA), PM, and lysosome toward the peroxisome/endosome. Colors correspond to spatial maps for AC16 cells throughout the manuscript. (Right) Ultracentrifugation fraction profile of DNAJB11, DNAJC3, DNAJC10, PDIA6, EMC4, EMC8, VAPA, and VAPB showing the localization of the proteins to the ER and to the peroxisome/endosome fraction in normal and thapsigargin-treated AC16 cells, respectively. X-axis: fraction 1 to 10 of ultracentrifugation. Y-axis: relative channel abundance. Bold lines represent the protein of interest; light lines represent ultracentrifugation profiles of all proteins classified to a respective localization. Colors correspond to subcellular localization in panel B and for all AC16 data throughout the manuscript. Numbers in the box represent BANDLE localization probability to the compartment. RNA Granule Score: score from RNA Granule Database (https://rnagranuledb.lunenfeld.ca/). A score of 7 or above is considered a known stress granule protein. Phi: predicted phase separation participation. Circle denotes a prediction of True within the database, X denotes a prediction of False. RBP: Annotated RNA binding protein on the RNA Granule Database. One circle denotes known RNA binding proteins (RBP) in at least one data set; two circles denote known RBP in multiple datasets. Dashes indicate proteins not found within the RNA Granule Database.

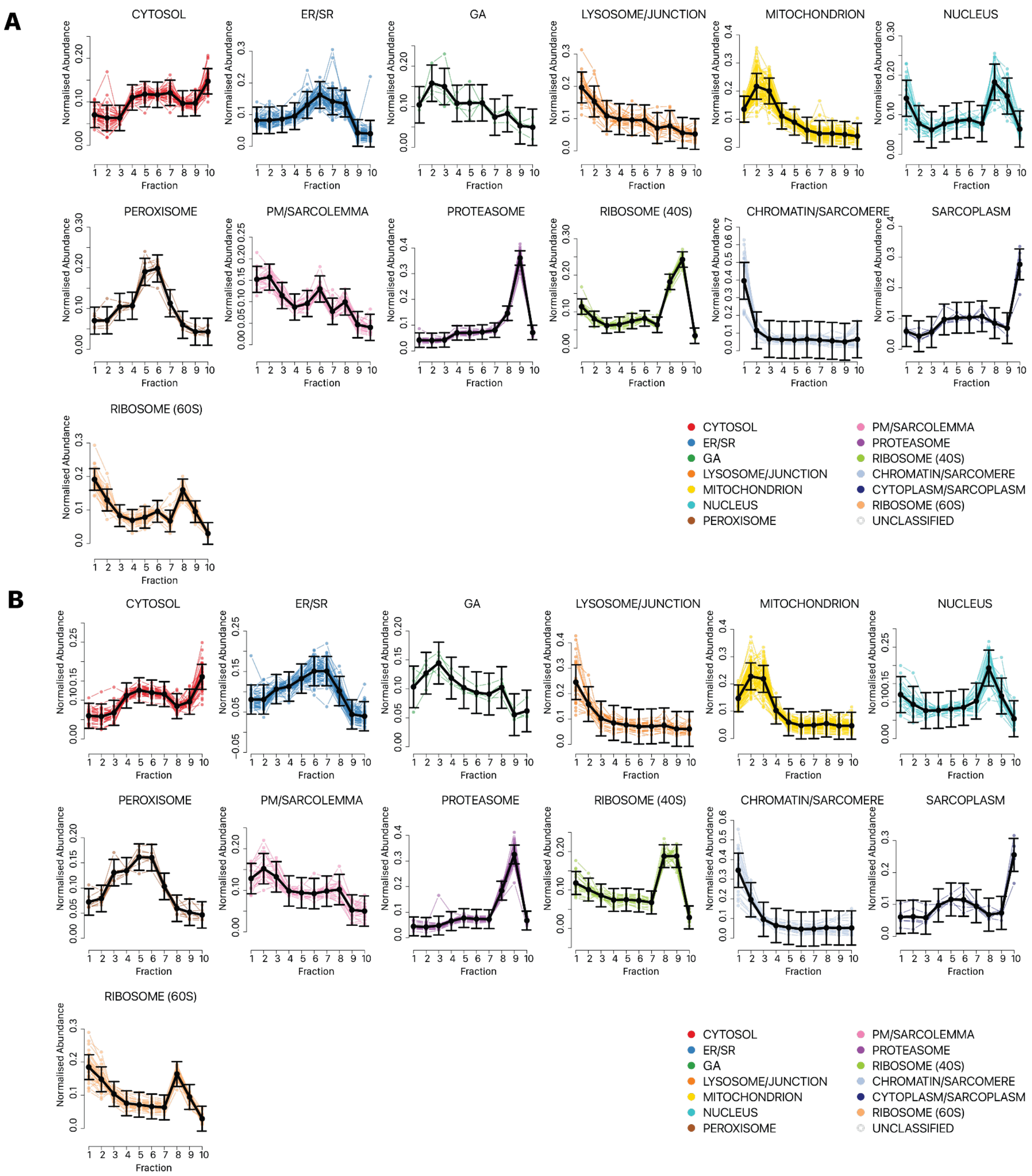


**Supplemental Figure S13.** Ultracentrifugation fraction distributions of cellular component markers in human iPSC-CMs. Additional iPSC-CM specific compartments and markers were curated manually, including a sarcomere and a cell junction compartment. The compartments were merged with the chromatin and the lysosome compartments due to similarity in sedimentation profile under the present ultracentrifugation scheme. Replicate one of each experimental condition is shown. **A.** Control iPSC-CM. **B.** iPSC-CM treated with 0.5 μM carfilzomib, 48 hours. The line plots show the normalized abundance (y-axis) of marker proteins for each subcellular localization experiment across ultracentrifugation fractions (x-axis) as measured by the TMT channel intensities. The fractions correspond to the ultracentrifugation steps in Supplemental Table S2. Colors correspond to spatial maps for iPSC-CMs cells throughout the manuscript. Black lines show average trend line and standard deviation, showing consistent sedimentation profiles upon carfilzomib treatment.

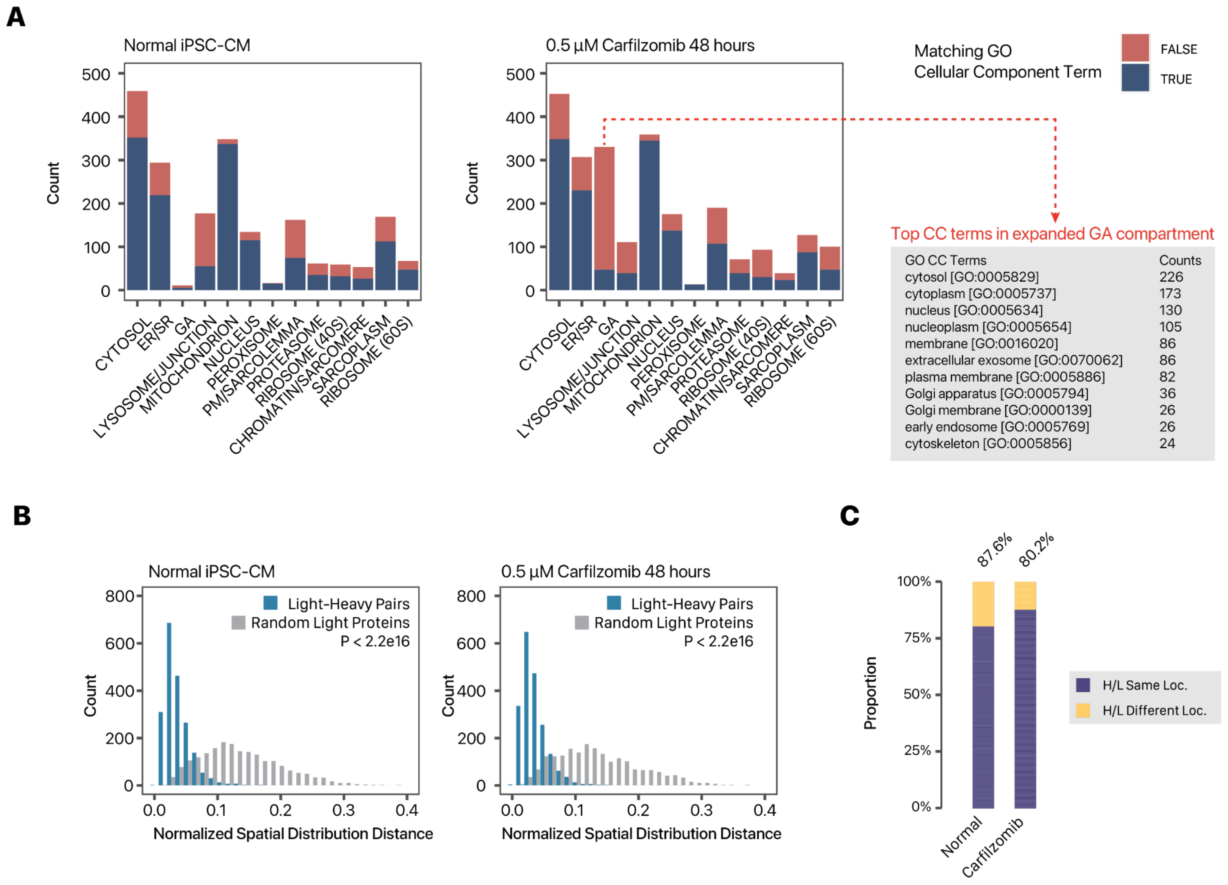


**Supplemental Figure S14.** Correspondence of spatial classification with prior annotations in human iPSC-CMs. **A.** The bar charts show the number of light (i.e., non-heavy-SILAC labeled) proteins (y-axis) classified to each of 13 subcellular locations (x-axis) in normal (left) and carfilzomib-treated (right) treated iPSC-CMs. Colors represent whether proteins classified to each subcellular location are also known to reside in the subcellular component of question in Gene Ontology Cellular Component terms retrieved from UniProt. In carfilzomib treated cells, a number of proteins are classified as co-sedimenting with Golgi markers; the top associated GO Cellular Component terms of these proteins are shown on the right and suggest they contain cytoplasmic proteins and proteins with multiple locations. In normal and carfilzomib-stressed cells, 70.8% and 63.0% of classified proteins are consistent with known annotations, respectively. **B.** Histogram showing the similarity in light and heavy proteins in normalized fraction abundance profiles in (left) normal and (right) carfilzomib-treated iPSC-CMs. X-axis: euclidean distance of fraction profiles across 2 replicates; y-axis: count. Blue: euclidean distance for quantified light-heavy protein pairs. Grey: distance of each corresponding light protein with a random sampled light protein. P value: Mann-Whitney test. **C.** In baseline and stressed iPSC-CMs, 87.6% and 80.2% of light and heavy protein pairs are assigned to the same subcellular localization.

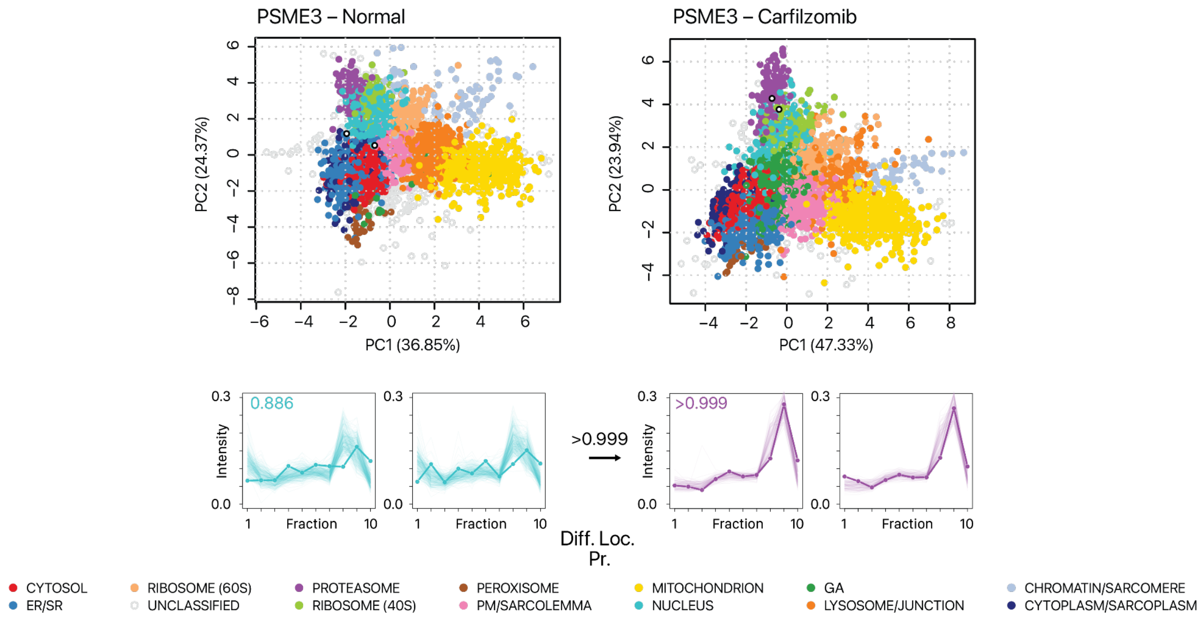


**Supplemental Figure S15.** Translocation of PA28/PSME3 upon carfilzomib. Spatial map (PC1 vs. PC2) and ultracentrifugation fraction profiles of PA28/PSME3 in normal and carfilzomib-treated human iPSC-CMs, showing a likely differential localisation from the nuclear to the proteasome compartment. Open circles: light and heavy PSME3 in each plot. Numbers inside the fraction profile correspond to BANDLE localization probability. Numbers at arrows correspond to BANDLE differential localization probability (Diff. Loc. Pr.).

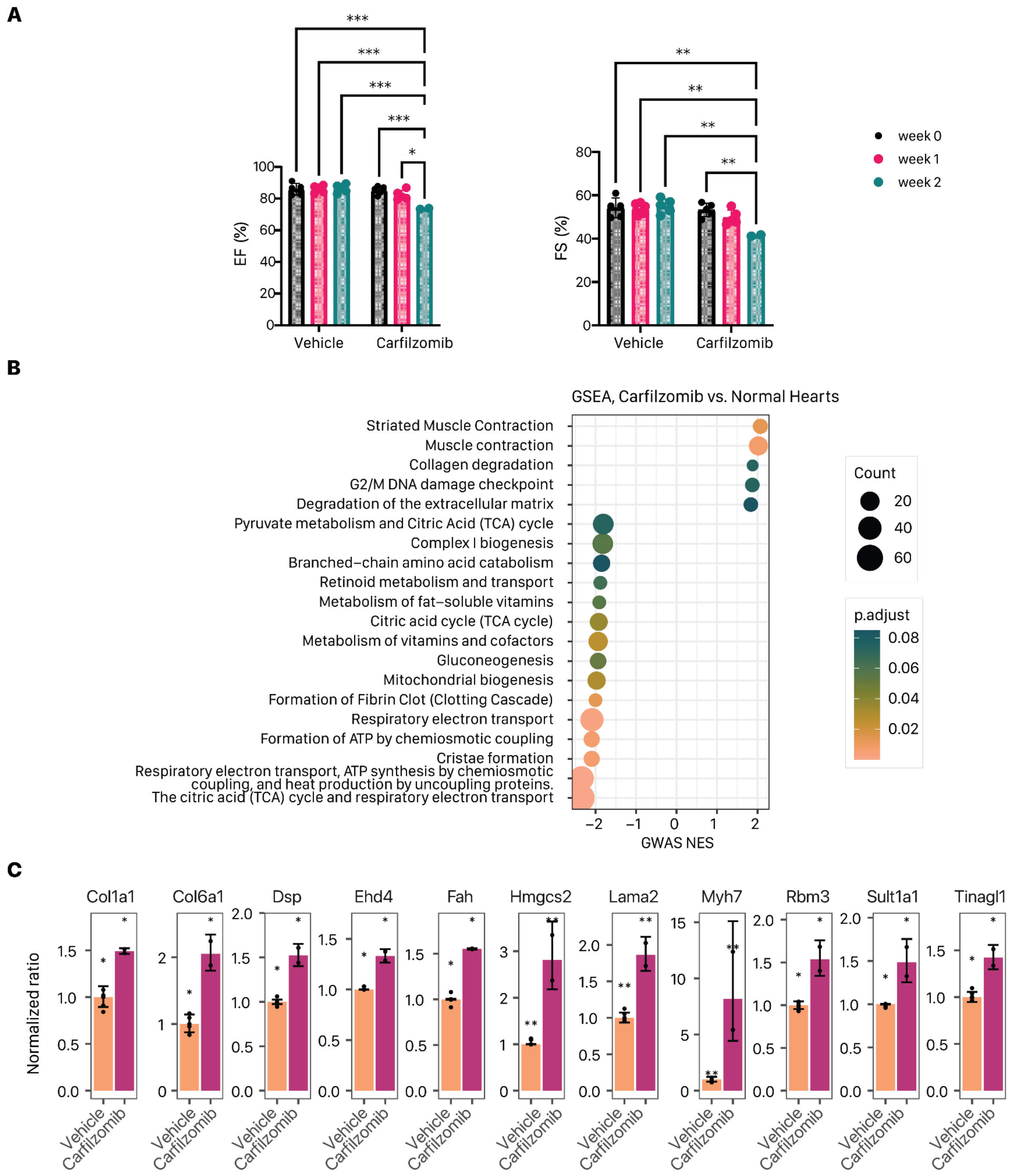


**Supplemental Figure S16.** In vivo cardiac effect of carfilzomib. **A.** Ejection fraction (EF) and fractional shortening (FS) of carfilzomib (CFZ) treated mice (n=5 for week 0 and week 1, n = 2 for week 2) and vehicle (Veh) treated mice (n = 5), injected twice weekly. Two-way ANOVA with Tukey's correction, p-value 0.0001 to 0.001: ***, p-value 0.001 to 0.01: **, p-value 0.01 to 0.05: * **B.** Gene set enrichment analysis of protein quantification (Carfilzomib vs. DMSO) showing a number of significantly enriched terms (y-axis) implicated in cardiac dysfunction and mitochondrial changes. Size denotes number of quantified proteins in the gene set, color: GSEA

FDR adjusted P value; x-axis: GSEA normalized enrichment score (NES). The suppression of mitochondrial proteins in vivo is consistent with the measured mitochondrial dysfunction in iPSC-CM treated with carfilzomib **C.** Two weeks of carfilzomib treatment led to 11 differentially expressed proteins at 5% FDR (25 at 10% FDR) in the mouse heart (n=2 for carfilzomib treated mice, n=5 for vehicle) out of 3379 quantified proteins. The significant proteins are visualized in bar charts to show the normalized expression in vehicle and carfilzomib treatment, highlighting the accumulation of MYH7 and DSP in carfilzomib. Error bars: median absolute deviation. *: limma FDR adjusted P < 0.05; **: limma FDR adjusted P < 0.01.
